# Loss of Arginase 2 Disrupts Striatum-Specific Polyamine Homeostasis

**DOI:** 10.1101/2025.06.02.657391

**Authors:** Martyna Nalepa, Omar Basheer, Mariusz Radkiewicz, Katarzyna Skowrońska, Aleksandra Skweres, Aleksandra Owczarek, Agata Dalka, Wojciech Hilgier, Magdalena Zielińska, Emilia Samborowska, Michał Węgrzynowicz

**Affiliations:** Laboratory of Molecular Basis of Neurodegeneration, Mossakowski Medical Research Institute, Polish Academy of Sciences, Pawińskiego 5, 02-106 Warsaw, Poland; Laboratory of Mass Spectrometry, Institute of Biochemistry and Biophysics, Polish Academy of Sciences, Pawińskiego 5a, 02-106 Warsaw, Poland; Department of Neurotoxicology, Mossakowski Medical Research Institute, Polish Academy of Sciences, Pawińskiego 5, 02-106 Warsaw, Poland; Doctoral School of Translational Medicine, Centre of Postgraduate Medical Education, Marymoncka 99/103, 01-813 Warsaw, Poland

**Keywords:** striatum, arginase 2, arginine, ornithine, polyamines

## Abstract

Arginase converts arginine (Arg) to ornithine (Orn), regulating their availability for the metabolic pathways that utilize these amino acids. The roles of arginase isoenzymes, Arg1 and Arg2, vary by cell type, tissue, and physiological state. In the brain, Arg2 is the predominant isoenzyme, particularly enriched in the striatum, where it localizes to a striatum-specific neuronal population - medium spiny neurons (MSNs). While the precise role of Arg2 in MSNs remains unclear, its loss alters the striatal metabolomic profile, highlighting its metabolic significance.

Here, to investigate the basis of these complex metabolic changes, we examined Arg metabolism in Arg2 knockout (Arg2^-/-^) mice. Targeted analysis of Arg-related metabolites and selected proteins regulating Arg metabolic pathways revealed that Arg2 loss significantly increased Arg levels but did not affect Orn, likely due to compensatory synthesis of Orn from Arg (via arginine:glycine amidinotransferase) and/or proline (via ornithine aminotransferase). Additionally, markers of nitric oxide (NO) production remained unchanged, suggesting that striatal Arg2 is not involved in the regulation of this pathway, a role commonly attributed to arginase. Most notably, Arg2 loss disrupted polyamine homeostasis, shifting the balance toward higher polyamines at the expense of lower ones and altering the expression of polyamine-regulating proteins. These findings highlight Arg2 crucial role in striatal metabolism and its potential relevance to striatum-related disorders. Given that striatal Arg2 impairment has been reported in Huntington’s disease, a neurodegenerative disorder specifically affecting MSNs, understanding its function may provide insights into the pathology.

## Introduction

Arginase is a manganese-dependent metalloenzyme that hydrolyzes arginine (Arg) into ornithine (Orn) and urea (Fig. 1) [1]. Arg is essential for multiple metabolic pathways and serves as a precursor for key metabolites like nitric oxide (NO), a signalling molecule, and creatine, important for energy metabolism. Orn can be used for the synthesis of proline (Pro), crucial for collagen production, glutamate (Glu), a major excitatory neurotransmitter, and polyamines – putrescine (Put), spermidine (Spmd), and spermine (Spm) – invoved in important cellular pathways like cell growth and differentiation and stress responses (Fig. 1) [2]. Arginase thus plays a central role in amine metabolism and critical cellular processes [1].

**Fig. 1.**
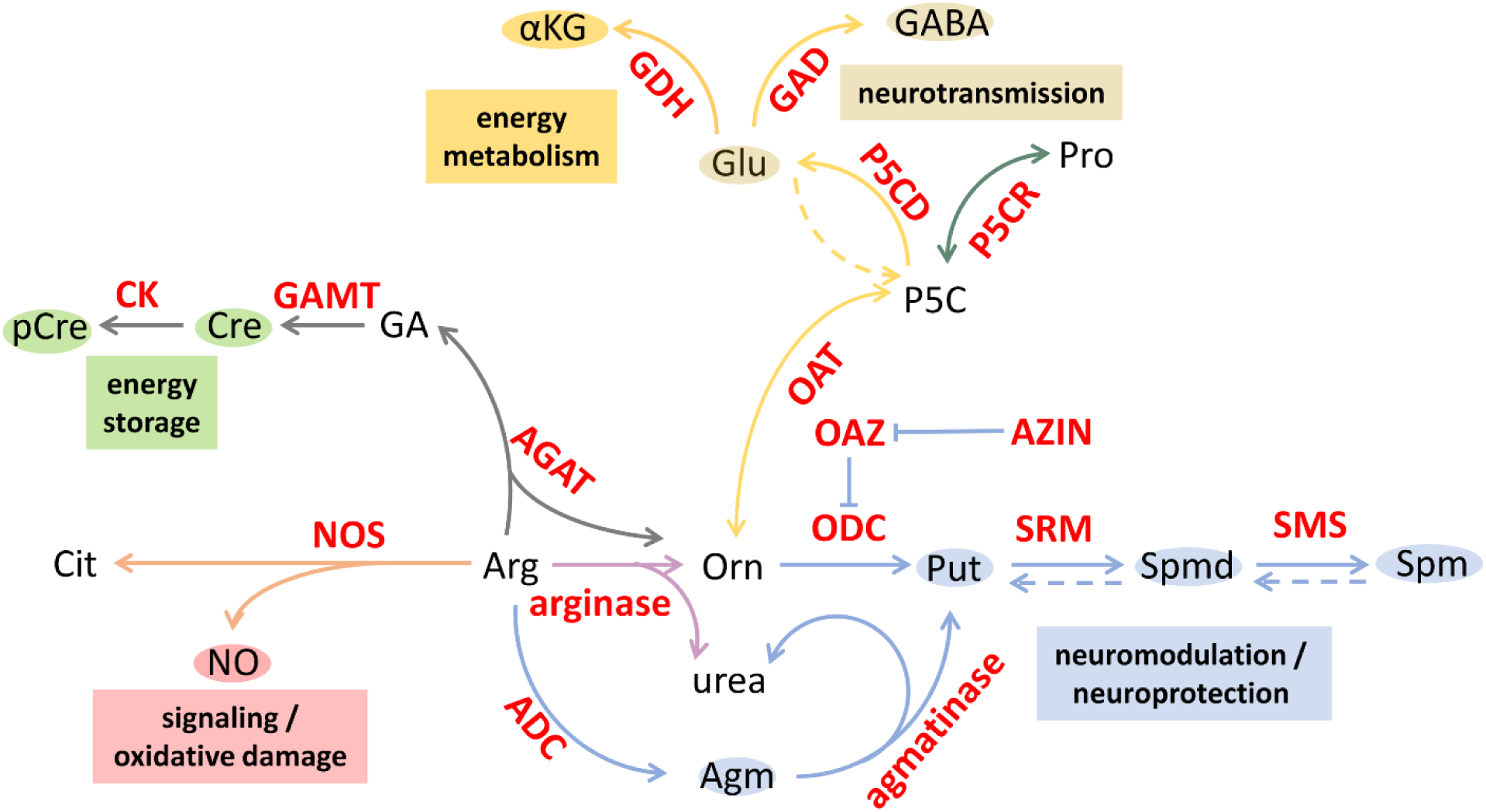
Schematic representation of arginine metabolism in the brain. αKG –α-ketoglutarate; **ADC** – arginine decarboxylase; **AGAT** – arginine:glycine amidinotransferase; **Agm** – agmatine; **Arg** – arginine; **AZIN** – antizime ihibitor; **Cit** – citrulline; **CK** – creatine kinase; **Cre** – creatine; **GA** – guanidinoacetate; **GABA** – γ-aminobutyric acid; **GAD** – glutamate decarboxylase; **GAMT** – guanidinoacetate methyltransferase; **GDH** – glutamate dehydrogenase; **Glu** – glutamate; **NO** – nitric oxide; **NOS** – nitric oxide synthase; **OAT** – ornithine aminotransferase; **OAZ –** ornithine decarboxylase antizyme; **ODC** – ornithine decarboxylase; **Orn** – ornithine; **P5C** – pyrroline-5-carboxylate; **P5CD –** pyrroline-5-carboxylate dehydrogenase; **P5CR** – pyrroline-5-carboxylate reductase; **pCre** – phosphocreatine; **Pro** – proline; **Put** – putrescine; **SMS** – spermine synthase; **Spm** – spermine; **Spmd** – spermidine; **SRM** – spermidine synthase.

Mammals express two isoenzymes: arginase 1 (Arg1) and arginase 2 (Arg2) [1]. Cytosolic Arg1 is highly expressed in the liver and is critical for the urea cycle, playing a crucial role in protecting from ammonia toxicity. Ammonia is especially toxic to the brain, and inborn Arg1 deficiency, if not properly managed, can lead to severe neurological impairments, including developmental delays, cognitive deficits, and, ultimately, death [3–5]. Cerebral Arg1 is well described for its role in glial cells, especially microglia, where it supports polarization towards the anti-inflammatory M2 phenotype. In non-inflammatory conditions, Arg1 is largely absent in most brain regions, suggesting a role as immune response modulator rather than baseline function [6].

Arg2 is predominantly a mitochondrial enzyme, and is less well characterized than Arg1. It regulates Arg and Orn levels across tissues influencing the availability of these compounds for the processes like protein synthesis, cell signalling, cellular growth or apoptosis. This isoenzyme therefore appears to contribute to precise regulation of cellular metabolism across various tissues. Arg2 upregulation has been shown to be involved in various pathological conditions, including atherosclerosis [7], pulmonary hypertension [8], erectile dysfunctions [9] or diabetic nephropathy [10]. In these conditions, mechanisms of Arg2 involvement in disease progression are complex and condition- and tissue-specific.

The exact role of Arg2 in the central nervous system remains unclear. One brain region of growing interest is the striatum, where Arg2 is the only arginase isoenzyme, and where it is exceptionally enriched. The striatum regulates motor functions and reward processing, and its dysfunction is implicated in certain neurodegenerative disorders, including Parkinson’s disease (PD), and, especially, Huntington’s disease (HD) [11]. HD is a genetic disorder characterized by involuntary movements and related with a degeneration of striatum-specific neuronal population – GABAergic medium spiny neurons (MSNs). An early and progressive deficit of Arg2 has been observed in HD mouse models [12], but its role in disease progression remains unclear. Recently, we found that Arg2 is specifically expressed in MSNs and that its loss disrupts striatal metabolic profiles, as measured by nuclear magnetic resonance (NMR) spectroscopy, highlighting the importance of Arg2 for striatal metabolism, and, consequently, for functioning of this region [13].

Here, we examined how Arg2 deficiency affects striatal Arg-related metabolic pathways. We compared Arg and its metabolites in the striata of wild-type C57 (Arg2^+/+^) and Arg2 knockout (Arg2^-/-^) mice and assessed the expression and distribution of key regulatory proteins. We found that Arg2 loss resulted in significant striatal Arg accumulation without changes in Orn, suggesting activation of alternative pathways (possibly through AGAT and OAT enzymes) maintaining Orn homeostasis independently of Arg2. Notably, NO production remained unchanged, suggesting that Arg2 doesn’t control striatal NO. Despite unchanged total polyamine levels, Arg2 loss significantly disrupted their balance by shifting it towards higher polyamines possibly related with altered expression of polyamine-resulted proteins including SRM, ODC, and AZIN1. PCA revealed distinct Arg-related metabolic profiles between Arg2^+/+^ and Arg2^-/-^ striata, indicating that Arg2 is crucial for striatum-specific Arg metabolism and polyamines homeostasis.

## Results

To explore metabolic role of Arg2 in the striatum, we performed a comparative analysis of the striata of Arg2^+/+^ and Arg2^-/-^ mice, the latter being efficiently depleted of striatal Arg2, as demonstrated in our recent report [13]. We measured metabolites related to Arg and Orn, Arg2 substrate and product, respectively (Fig. 2A), and assessed expression of enzymes involved in Arg-related pathways. Selected proteins were also examined by immunohistochemistry to support the metabolic data.

**Fig. 2.**
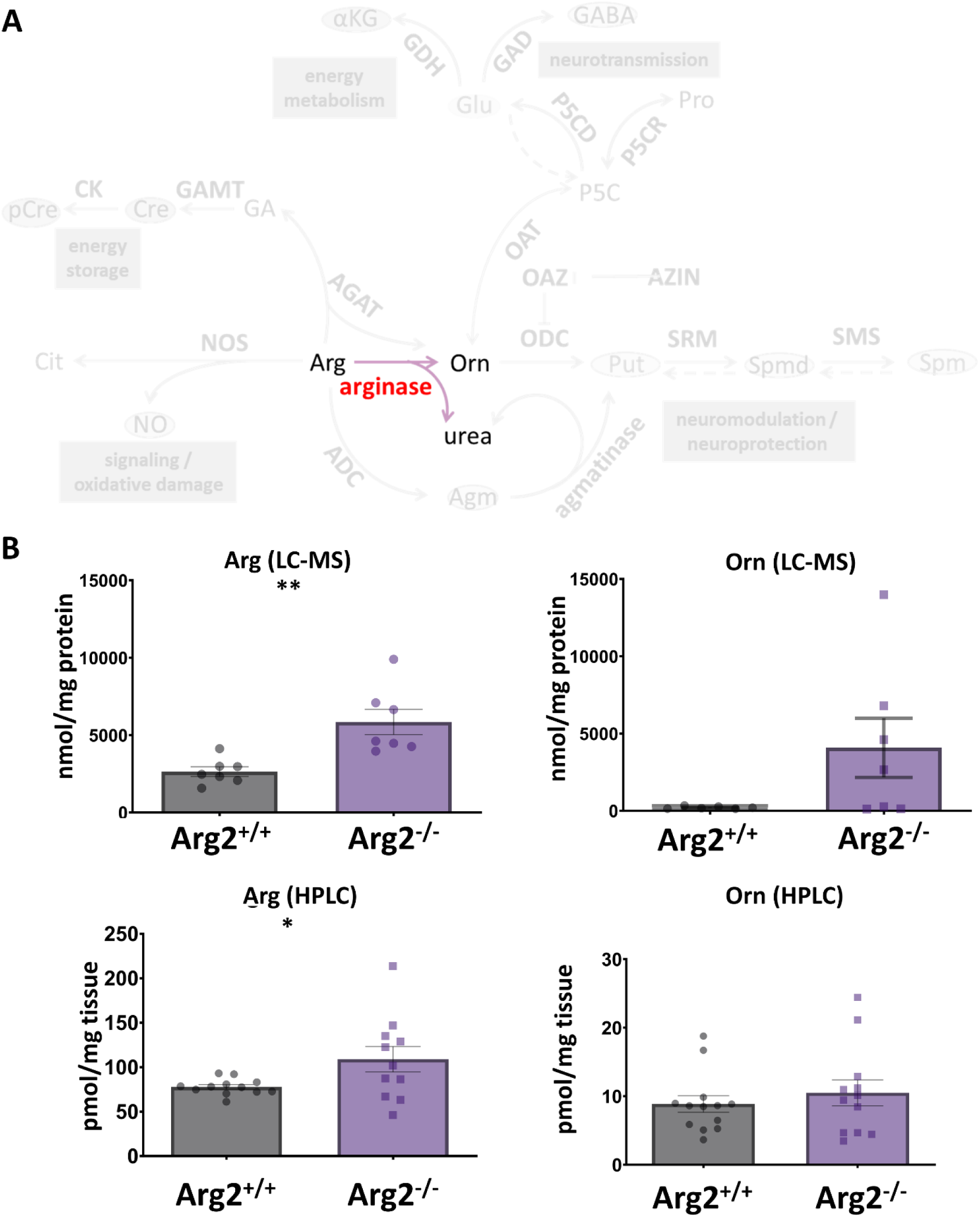
Analysis of arginine (Arg) and ornithine (Orn), substrate and product of arginase in the striatum of Arg2^+/+^ and Arg2^-/-^ mice. **(A)** Schematic representation of arginase-catalyzed conversion of Arg to Orn and urea. **(B)** Measurements of Arg and Orn content in Arg2^+/+^ and Arg2^-/-^ mouse striata. (Top left panel) A statistically significant decrease in Arg levels was observed in Arg2^-/-^ striatum as measured by LC-MS (2644.34 ± 308.89 nmol/mg protein in Arg2^+/+^ vs 5849.70 ± 818.26 nmol/mg protein in Arg2^-/-^; p = 0.003), n = 7. (Top right panel) Despite the apparent trend, no statistically significant differences in the levels of Orn were detected between Arg2^+/+^ and Arg2^-/-^ striata as measured by LC-MS (209.79 ± 33.30 nmol/mg protein in Arg2^+/+^ vs 4079.80 ± 1913.99 nmol/mg protein in Arg2^-/-^; p = 0.090), n = 7. (Bottom left panel) A statistically significant increase in Arg levels was observed in Arg2^-/-^ striatum as measured by HPLC (77.87 ± 2.58 pmol/mg tissue in Arg2^+/+^ vs 109.02 ± 14.30 pmol/mg tissue in Arg2^-/-^; p = 0.036), n = 10-12. (Bottom right panel) No statistically significant differences in Orn levels were observed between Arg2^+/+^ and Arg2^-/-^ striata as measured by HPLC (8.86 ± 1.21 pmol/mg tissue in Arg2^+/+^ vs 11.01 ± 1.99 pmol/mg tissue in Arg2^-/-^; p = 0.350), n = 11-13. Data are shown as mean ± SEM. *p < 0.05, **p < 0.01, t-test.

### Arg2 loss results in striatal Arg accumulation without affecting Orn

As expected, Arg2 deletion resulted in a significant accumulation of Arg in the striatum, as shown by LC-MS (Fig. 2B). Surprisingly, Orn levels didn’t differ significantly between Arg2^+/+^ and Arg2^-/-^ mice, although Arg2^-/-^ mice showed high variability and a trend towards increase (Fig. 2B). These findings were confirmed by HPLC in an independent cohort, again showing increased Arg levels in Arg2^-/-^ striata (Fig. 2B), with no significant differences in Orn (Fig. 2B). HPLC-measured Orn levels were more consistent across individuals compared to LC-MS data.

### Arg2 is not involved in controlling NO synthesis in the striatum

To investigate whether Arg2 regulates striatal NO synthesis by competing with nitric oxide synthases (NOS) for Arg (Fig. 3A), we measured nitrates and nitrites, collectively referred to as NOx, as indicators of NO production. We found no differences in striatal NOx levels between Arg2^+/+^ and Arg2^-/-^ mice (Fig. 3B). We also measured citrulline (Cit), the co-product of Arg-to-NO conversion by NOS (Fig. 3A), and, again, observed no effect of Arg2 loss as measured by both, LC-MS and HPLC (Fig. 3B). These data indicate that Arg2 doesn’t influence NO production in the striatum and excess Arg due to Arg2 deficiency is not utilized by NOS. Additionally, we found that the levels of nNOS (neuronal NOS isoform) remained unchanged (Fig. 3C), demonstrating no compensatory response of nNOS to Arg2 deficiency.

**Fig. 3.**
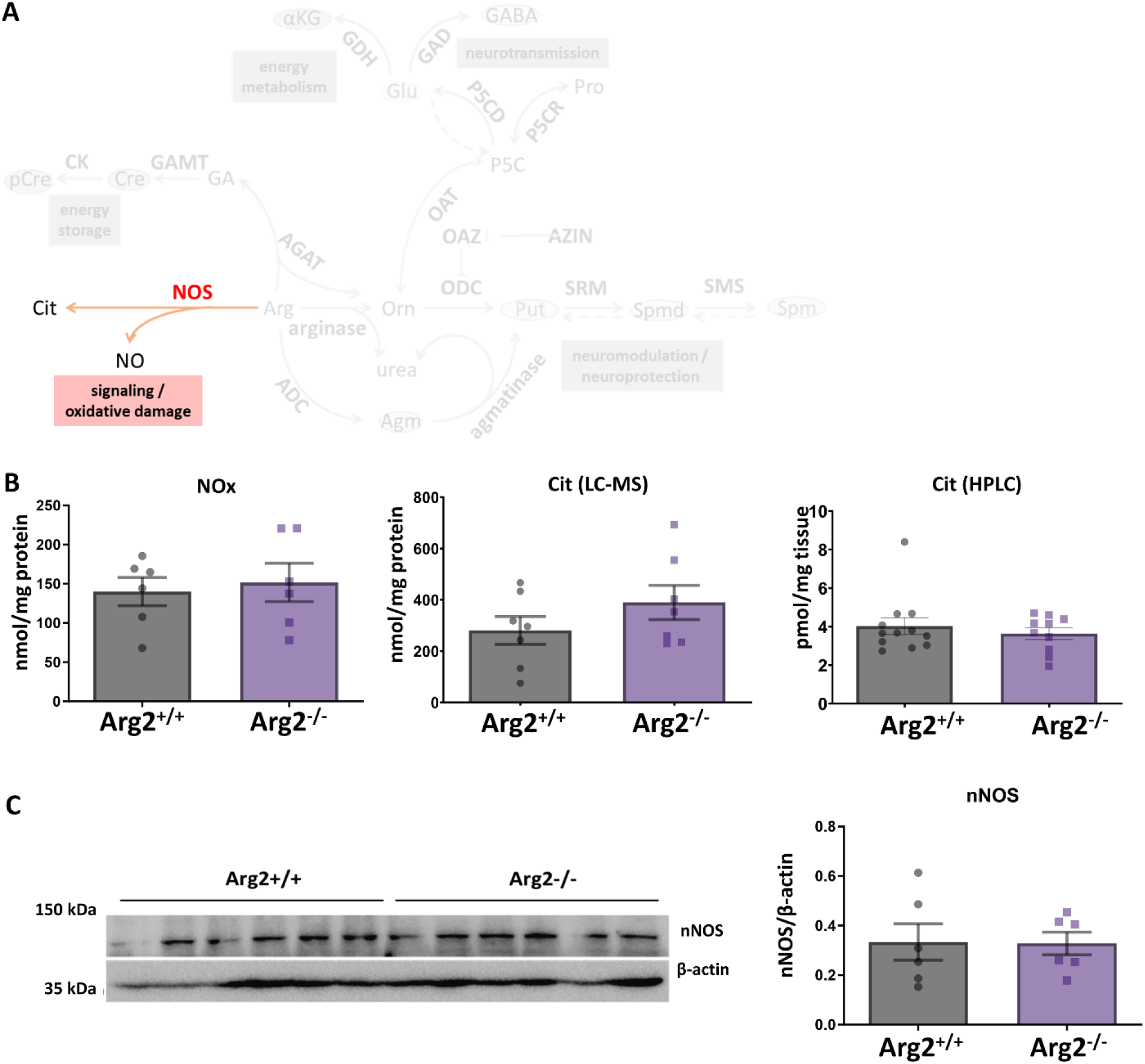
Analysis of nitric oxide (NO)-related markers in the striatum of Arg2^+/+^ and Arg2^-/-^ mice. **(A)** Schematic representation of NO synthesis pathway. **(B)** Measurements of NOx (combined levels of nitrates and nitrites) and citrulline (Cit) content in Arg2^+/+^ and Arg2^-/-^ mouse striata. (Left panel) No statistically significant differences in NOx were observed between Arg2^+/+^ and Arg2^-/-^ striata (140.02 ± 18.06 nmol/mg of protein in Arg2^+/+^ vs 151.78 ± 24.33 nmol/mg of protein in Arg2^-/-^; p = 0.706), n = 6. (Middle panel) No statistically significant differences in Cit were observed between Arg2^+/+^ and Arg2^-/-^ striata as measured by LC-MS (280.68 ± 54.57 nmol/mg protein in Arg2^+/+^ vs. 389.71 ± 66.80 in Arg2^-/-^; p = 0.230), n = 7. (Right panel) No statistically significant differences in Cit levels were observed between Arg2^+/+^ and Arg2^-/-^ striata as measured by HPLC (4.03 ± 0.44 pmol/mg tissue in Arg2^+/+^ vs 3.63 ± 0.30 in Arg2^-/-^; p = 0.484), n = 10-12. **(C)** Measurements of neuronal NO synthase (nNOS) expression in Arg2^+/+^ and Arg2^-/-^ mouse striata. (Left panel) Western blot image of nNOS and β-actin in Arg2^+/+^ and Arg2^-/-^ striata. (Right panel) No statistically significant differences in nNOS expression were observed between Arg2^+/+^ and Arg2^-/-^ striata as determined by the quantification of nNOS western blot normalized to β-actin western blot (0.334 ± 0.074 in Arg2^+/+^ vs 0.328 ± 0.045 in Arg2^-/-^ (arbitrary units); p = 0.947), n = 6. Data are shown as mean ± SEM.

### Arg2 loss induces compensatory mechanisms to preserve striatal Orn levels: possible roles of AGAT and OAT

Unchanged striatal Orn levels in Arg2^-/-^ mice (Fig. 2B) suggests compensatory mechanisms maintaining Orn homeostasis. Our earlier NMR-based metabolomics analysis identified Cre pathway metabolites – guanidinoacetate (GA), Cre and creatinine as key contributors to differences between Arg2^+/+^ and Arg2^-/-^ striata [13]. GA, increased in Arg2^-/-^ striata, is produced by arginine:glycine amidinotransferase (AGAT) from Arg and Gly with Orn as co-product (Fig. 4A), suggesting AGAT upregulation as an alternative pathway for Arg-to-Orn conversion in Arg2 deficiency. We measured AGAT and indeed, we confirmed a significant increase in its expression in Arg2^-/-^ striata (Fig. 4B). Immunostaining showed AGAT localizing mainly in DARPP-32-negative neuropil, with stronger signal in Arg2^-/-^ striata (Fig. 4C). These results support GA increase in Arg2-deficient striata and may partially explain stable Orn levels despite absence of Arg2.

**Fig. 4.**
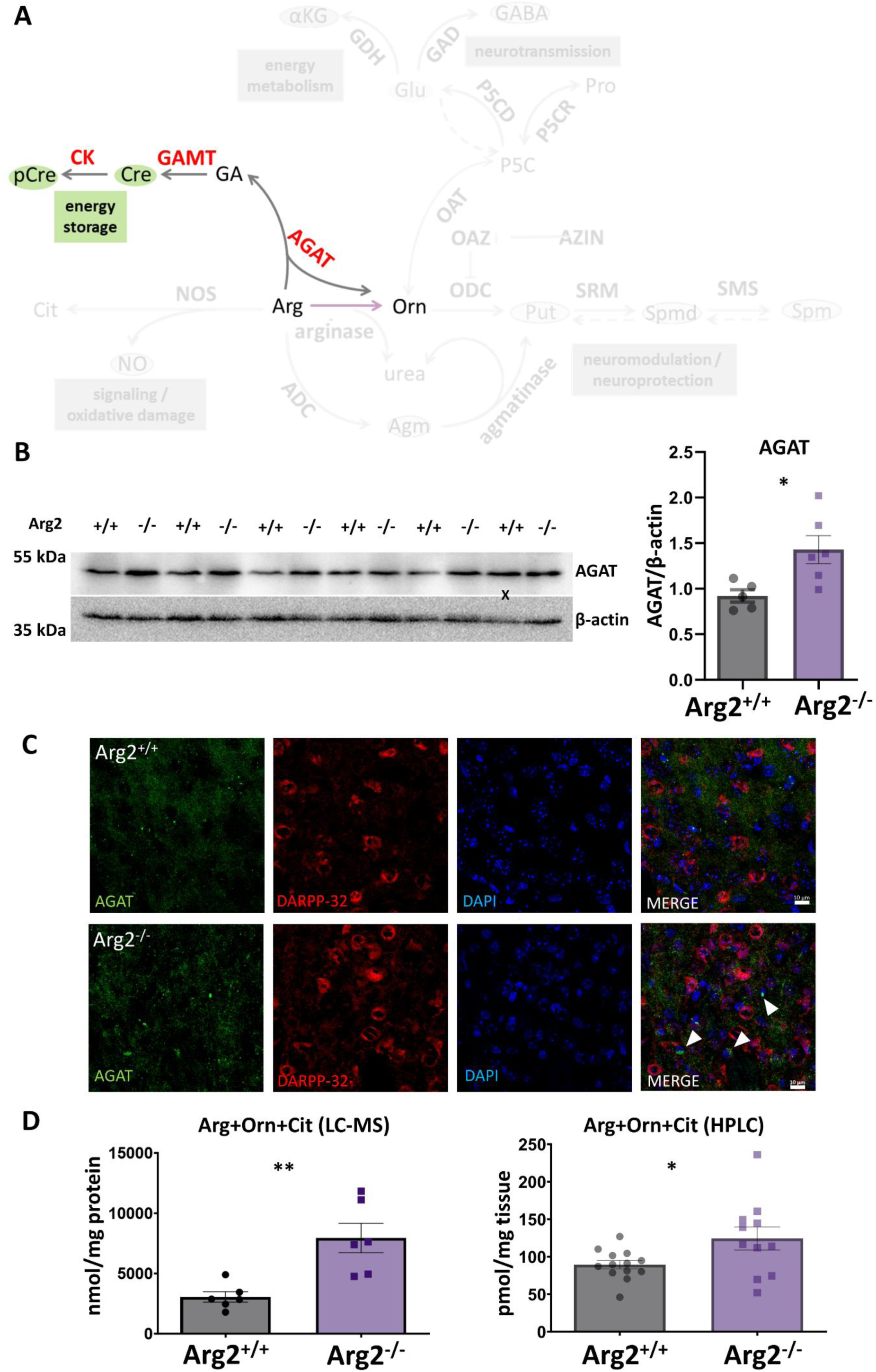
Analysis of arginine:glycine amidinotransferase (AGAT) expression as a potential mediator of alternative arginine-to-ornithine conversion pathway in the striatum of Arg2^-/-^ mice. **(A)** Schematic representation of creatine (Cre) synthesis pathway. **(B)** Measurements of AGAT expression in Arg2^+/+^ and Arg2^-/-^ mouse striata. (Left panel) Western blot image of AGAT and β-actin in Arg2^+/+^ and Arg2^-/-^ striata. (Right panel) A statistically significant increase in AGAT expression was observed in Arg2^-/-^ striata compared to Arg2^+/+^ striata as determined by the quantification of AGAT western blot normalized to β-actin western blot (0.92 ± 0.07 in Arg2^+/+^ vs 1.43 ± 0.15 in Arg2^-/-^ (arbitrary units); p = 0.019), n = 5-6. X indicates a sample identified as a technical outlier and excluded from the calculations. **(C)** Immunohistochemical analysis of AGAT distribution in Arg2^+/+^ and Arg2^-/-^ mouse striata. Co-immunofluorescent staining of AGAT (green) and the medium spiny neuron marker, DARPP-32 (red), in the striata of Arg2^+/+^ and Arg2^-/-^ mice. Nuclei are counterstained with DAPI (blue). Representative images for Arg2^+/+^ and Arg2^-/-^ are shown. Scale bars (10 μm) are located in the bottom right corner of the merged images. AGAT did not co-localize with DARPP-32. AGAT staining intensity was increased in Arg2^-/-^ striata, where accumulation of small, intense AGAT-positive clusters were found (arrows). **(D)** Analysis of combined levels of Arg and its direct amino acid products in the striatum of Arg2^+/+^ and Arg2^-/-^ mice. (Left panel) A statistically significant increase in the combined Arg+Cit+Orn levels was observed in the Arg2^-/-^ striatum, as measured by LC-MS (3050.55 ± 431.44 nmol/mg protein in Arg2^+/+^ vs 7942.03 ± 1219.22 nmol/mg protein in Arg2^-/-^; p = 0.004), n = 6. (Right panel) A statistically significant increase in the combined Arg+Cit+Orn levels was observed in the Arg2^-/-^ striatum, as measured by HPLC (89.37 ± 5.52 pmol/mg tissue in Arg2^+/+^ vs 124.35 ± 15.39 pmol/mg tissue in Arg2^-/-^; p = 0.032), n = 11-13. Data are shown as mean ± SEM. *p < 0.05, **p < 0.01, t-test.

Although Arg levels increased with Arg2 loss, Orn remained unchanged (Fig. 2B), suggesting that AGAT upregulation alone doesn’t fully account for the maintenance of Orn in Arg2^-/-^ striata. Total levels of Arg and its metabolites (Orn and Cit) were significantly elevated in Arg2^-/-^ striata, as shown by LC-MS and HPLC (Fig. 4D) suggesting involvement of additional pathway(s) of Orn production from non-Arg substrate.

Orn can be synthesized from Pro by ornithine aminotransferase (OAT) via the intermediate, 1-Pyrroline-5-carboxylic acid (P5C) (Fig. 5A). We measured Pro levels in striatal homogenates using LC-MS but found no significant differences between Arg2^+/+^ and Arg2^-/-^ mice (Fig. 5B). Combined Pro and Orn levels didn’t differ neither, but the Orn/Pro ratio was significantly higher in Arg2^-/-^ striata (Fig. 5B), suggesting a shift in Orn-Pro homeostasis towards Orn production. Supporting this, OAT protein expression was significantly increased in Arg2^-/-^ mice (Fig. 5C). Immunofluorescence analysis showed OAT primarily in DARPP-32-negative neuropil, with an increased signal in Arg2^-/-^ striata (Fig. 5D). These findings suggest enhanced Pro-to-Orn conversion as a compensatory mechanism alongside AGAT activation in response to Arg2 loss.

**Fig. 5.**
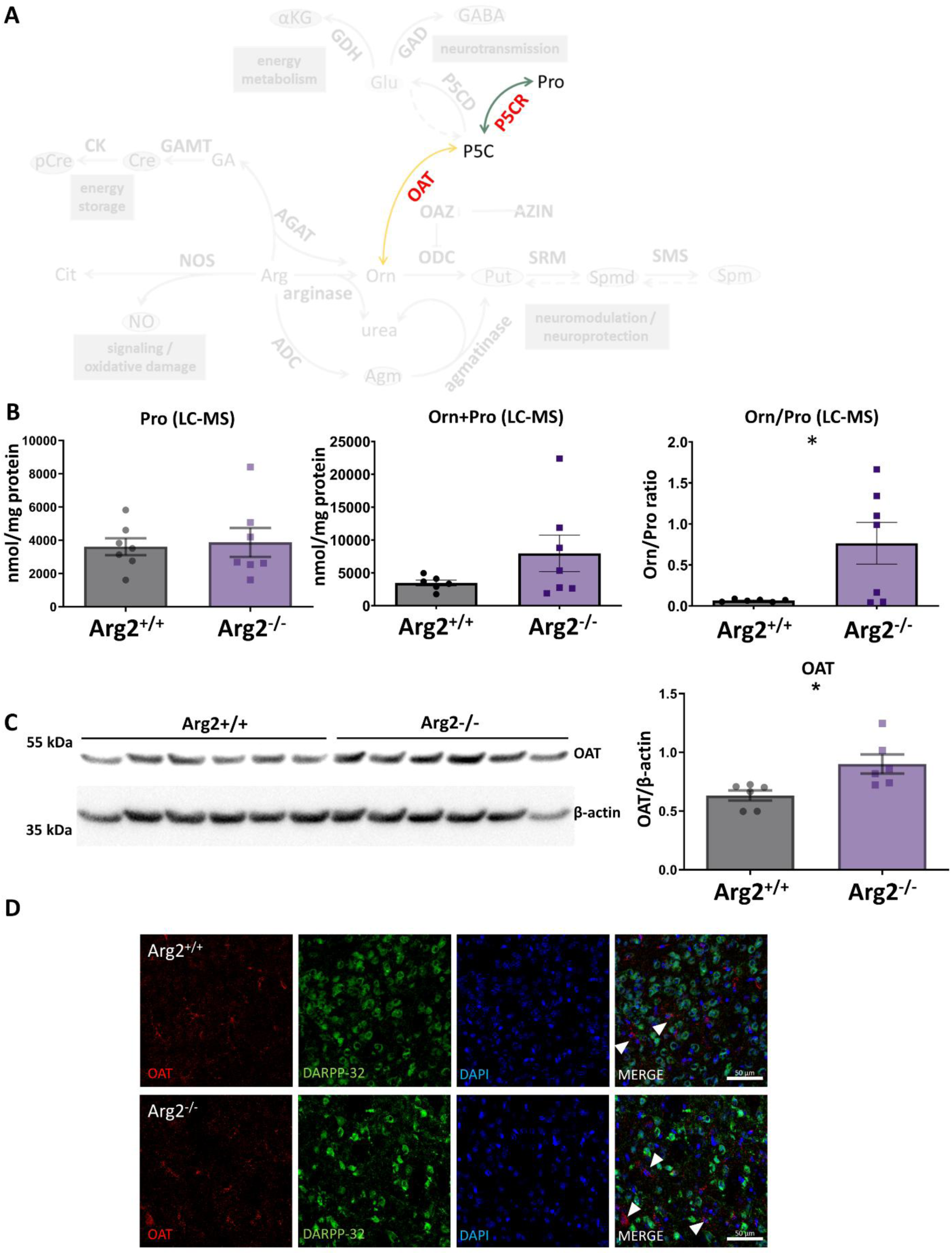
Analysis of proline (Pro) as a potential source of ornithine (Orn) via ornithine aminotransferase (OAT)-catalyzed pathway in the striatum of Arg2^-/-^ mice. **(A)** Schematic representation of bidirectional Pro to Orn interconversion pathway. **(B)** LC-MS-based measurements of Pro content in Arg2^+/+^ and Arg2^-/-^ mouse striata. (Left panel) No statistically significant differences in Pro levels were observed between Arg2^+/+^ and Arg2^-/-^ striata (3612 ± 508.30 nmol/mg protein in Arg2^+/+^ vs 3878.46 ± 871.20 nmol/mg protein in Arg2^-/-^; p = 0.796), n = 7. (Middle panel) No statistically significant differences in the combined Orn+Pro levels were observed between Arg2^+/+^ and Arg2^-/-^ striata (3454.48 ± 444.07 nmol/mg protein in Arg2^+/+^ vs 7958.26 ± 2774.96 nmol/mg protein in Arg2^-/-^; p = 0.167), n = 6-7. (Right panel) A statistically significant increase in the Orn/Pro ratio was observed in the Arg2^-/-^ striatum (0.066 ± 0.006 in Arg2^+/+^ vs 0.764 ± 0.254 in Arg2^-/-^; p = 0.028), n = 6-7. **(C)** Measurements of OAT expression in Arg2^+/+^ and Arg2^-/-^ mouse striata. (Left panel) Western blot image of OAT and β-actin in Arg2^+/+^ and Arg2^-/-^ striata. (Right panel) A statistically significant increase in OAT expression was observed in Arg2^-/-^ striata compared to Arg2^+/+^ striata as determined by the quantification of OAT western blot normalized to β-actin western blot (0.632 ± 0.044 in Arg2^+/+^ vs 0.901 ± 0.081 in Arg2^-/-^ (arbitrary units); p = 0.016), n = 6. **(D)** Immunohistochemical analysis of OAT distribution in Arg2^+/+^ and Arg2^-/-^ mouse striata. Co-immunofluorescent staining of OAT (red) and the medium spiny neuron marker, DARPP-32 (green), in the striata of Arg2^+/+^ and Arg2^-/-^ mice. Nuclei are counterstained with DAPI (blue). Representative images for Arg2^+/+^ and Arg2^-/-^ are shown. Scale bars (50 μm) are located in the right bottom corner of the merged images. OAT did not co-localize with DARPP-32, being primarily present in the DARPP-32-negative neuropil, and, sparsely in DARPP-32-negative neuronal bodies (arrows). Neuropil OAT staining intensity was increased in Arg2^-/-^ striatum. Data are shown as mean ± SEM. *p < 0.05, t-test.

Since Arg2 loss appears to direct the striatal Pro-P5C-Orn bidirectional pathway towards Orn synthesis (Fig. 5), we asked whether Glu, another P5C precursor (Fig. 6A) could also contribute to Orn recovery in Arg2^-/-^ striata. HPLC analysis, however, showed no changes in Glu or its direct metabolites, γ-aminobutyric acid (GABA) and glutamine (Gln) (Fig. 6B), nor in the Orn/Glu or Orn/(Glu+Gln+GABA) ratios (Fig. 6C). We also measured α-ketoglutarate (αKG), a Krebs cycle intermediate that can be produced from Glu by Glu dehydrogenase (Fig. 6A), and, similarly, found no differences between genotypes (Fig. 6D). These results suggests that Arg2 doesn’t significantly influence striatal Glu metabolism and Glu doesn’t contribute to Orn homeostasis in Arg2^-/-^ striata. Notably, aside from Arg, none of the 18 amino acids measured in our HPLC assay differed between Arg2^+/+^ and Arg2^-/-^ (Fig. 7).

**Fig. 6.**
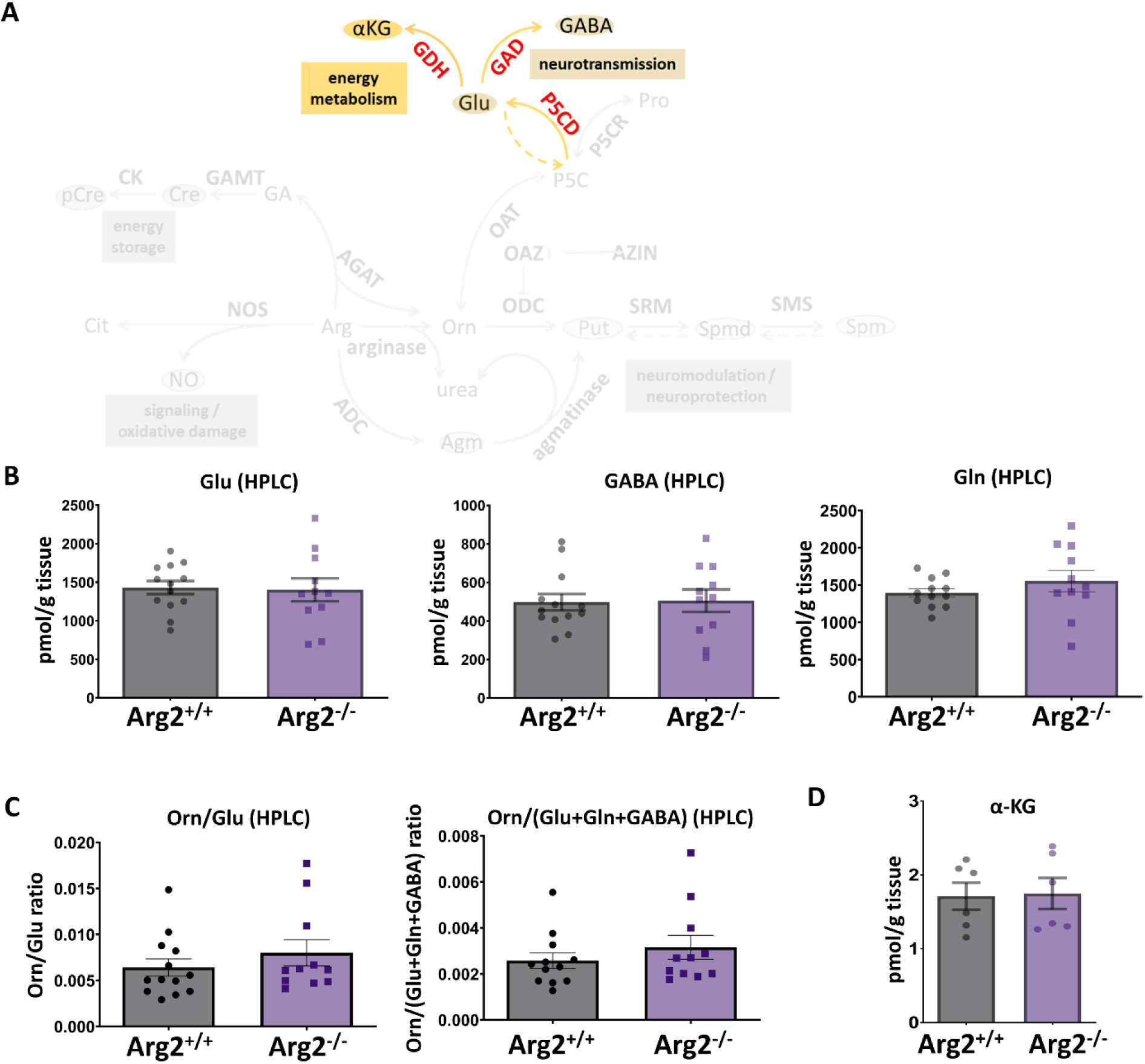
Analysis of glutamate (Glu) pathway in the striatum of Arg2^+/+^ and Arg2^-/-^ mice. **(A)** Schematic representation of the synthesis pathway of Glu and its derivatives from 1-Pyrroline-5-carboxylic acid (P5C). **(B)** HPLC-based measurements of Glu, γ-aminobutyric acid (GABA), and glutamine (Gln) content in Arg2^+/+^ and Arg2^-/-^ mouse striata. (Left panel) No statistically significant differences in Glu were observed between Arg2^+/+^ and Arg2^-/-^ striata (1429.67 ± 85.23 pmol/mg tissue in Arg2^+/+^ vs 1402.26 ± 148.06 pmol/mg tissue in Arg2^-/-^; p = 0.869), n = 11-13. (Middle panel) No statistically significant differences in GABA were observed between Arg2^+/+^ and Arg2^-/-^ striata (498.11 ± 42.56 pmol/mg tissue in Arg2^+/+^ vs 505.72 ± 58.04 pmol/mg tissue in Arg2^-/-^; p = 0.915), n = 11-13. (Right panel) No statistically significant differences in Gln were observed between Arg2^+/+^ and Arg2^-/-^ striata (1394.77 ± 56.97 pmol/mg tissue in Arg2^+/+^ vs 1553.61 ± 143.28 pmol/mg tissue in Arg2^-/-^; p = 0.299), n = 12-13. **(C)** HPLC-based measurements of the ratios of Orn to Glu and to combined Glu and its direct amino acid metabolites in Arg2^+/+^ and Arg2^-/-^ mouse striata. (Left panel) No statistically significant differences in the Orn/Glu ratio were observed between Arg2^+/+^ and Arg2^-/-^ striata (0.00641 ± 0.00093 in Arg2^+/+^ vs 0.00801 ± 0.0014 in Arg2^-/-^; p = 0.341), n = 11-13. (Right panel) No statistically significant differences in the Orn/(Glu+Gln+GABA) ratio were observed between Arg2^+/+^ and Arg2^-/-^ striata (0.00257 ± 0.00034 in Arg2^+/+^ vs 0.00315 ± 0.00052 in Arg2^-/-^; p = 0.356), n = 11-12. **(D)** Measurements of α-ketoglutarate (α-KG) content in Arg2^+/+^ and Arg2^-/-^ mouse striata. No statistically significant differences in α-KG were observed between Arg2^+/+^ and Arg2^-/-^ striata (1.71 ± 0.18 in Arg2^+/+^ vs 1.75 ± 0.21 in Arg2^-/-^; p = 0.900), n = 6. Data are shown as mean ± SEM.

**Fig. 7.**
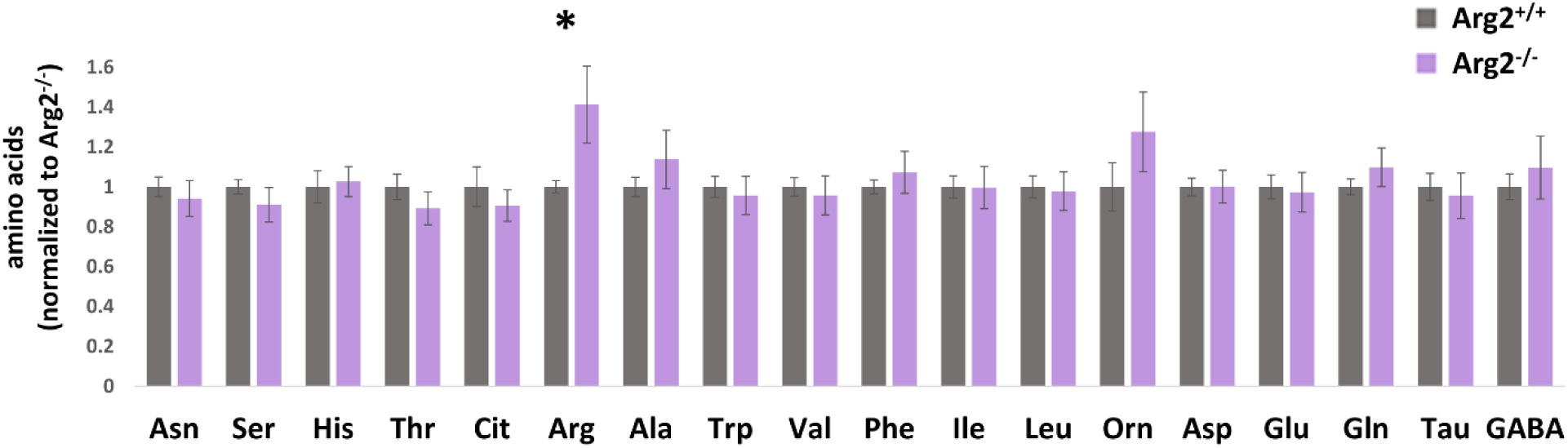
Analysis of amino acid profiles in the striatum of Arg2^+/+^ and Arg2^-/-^ mice. No statistically significant differences between Arg2^+/+^ and Arg2^-/-^ striata were observed for any amino acids except for Arg, as measured by HPLC. **Ala** – alanine; **Arg** – arginine; **Asn** – asparagine; **Asp** – aspartate; **Cit** – citrulline; **GABA** – γ-aminobutyric acid; **Gln** – glutamine; **Glu** – glutamine; **His** – histidine; **Ile** – isoleucine; **Leu** – leucine; **Orn** – ornithine; **Phe** – phenylalanine; **Ser** – serine; **Tau** – taurine; **Thr** – threonine; **Trp** – tryptophan; **Val** – valine. Data are shown as mean ± SEM normalized to individual amino acid control (mean Arg2^+/+^), n=10-13. *p < 0.05, t-test.

### Disrupted polyamine homeostasis in Arg2^-/-^ striatum

Orn is essential for polyamine synthesis, serving as a substrate for an initial, committed step of the pathway – Put synthesis by Orn decarboxylase (ODC). Put is then converted to Spmd by Spmd synthase (SRM) and Spmd to Spm by Spm synthase (SMS) (Fig. 8A). LC-MS analysis showed significantly reduced Put levels in Arg2^-/-^ striata, unchanged Spmd and increased Spm (Fig. 8B). Importantly, total polyamine content (Put+Spmd+Spm) remained unchanged (Fig. 8C), suggesting that Arg2 loss alters polyamine interconversion without affecting overall levels. This was reflected in increased ratios of higher-to-lower polyamines: Spmd/Put, Spm/Spmd, Spm/Put and (Spmd+Spm)/Put (Fig. 8C). We also measured polyamine degradation intermediates: N1-acetylspermine (N1-AcSpm) and N1-acetylspermidine (N1-AcSpmd). While N1-AcSpm was undetectable in both groups, N1-AcSpmd didn’t differ between Arg2^+/+^ and Arg2^-/-^ (Fig. 8D), suggesting altered Spmd-to-degradation doesn’t account for Put reduction in Arg2^-/-^ striata.

**Fig. 8.**
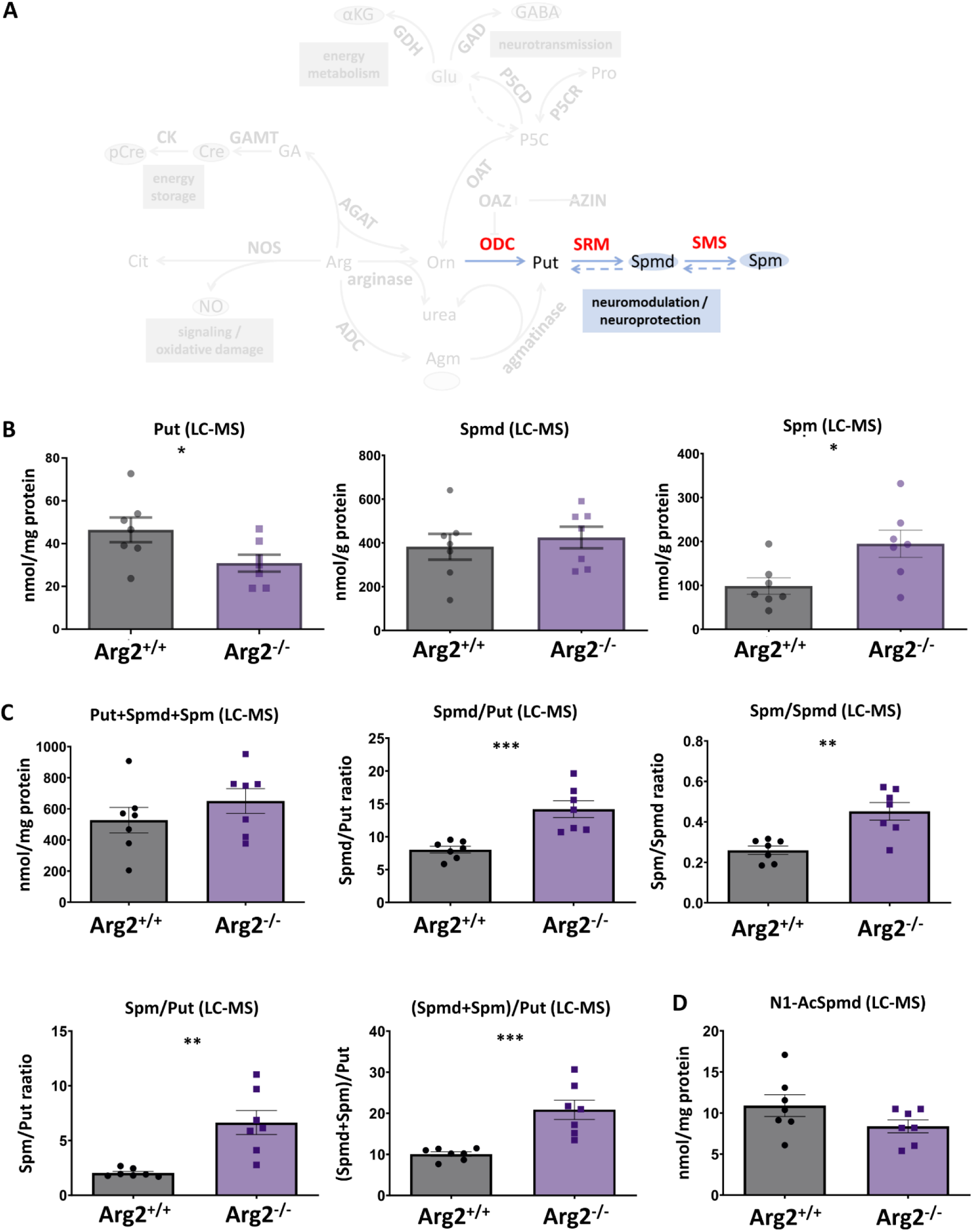
Analysis of polyamine pathway in the striatum of Arg2^+/+^ and Arg2^-/-^ mice. **(A)** Schematic representation of polyamine synthesis pathway. **(B)** LC-MS-based measurements of putrescine (Put), spermidine (Spmd) and spermine (Spm) content in Arg2^+/+^ and Arg2^-/-^ mouse striata. (Left panel) A statistically significant decrease in Put levels was observed in Arg2^-/-^ striatum (46.42 ± 5.79 nmol/mg protein in Arg2^+/+^ vs. 30.86 ± 3.99 nmol/mg protein in Arg2^-/-^; p = 0.047), n = 7. (Middle panel) No statistically significant differences in Spmd were observed between Arg2^+/+^ and Arg2^-/-^ striata 382.30 ± 59.10 nmol/mg protein in Arg2^+/+^ vs. 424.48 ± 49.41 nmol/mg protein in Arg2^-/-^; p = 0.594), n = 7. (Right panel) A statistically significant increase in Spm levels was observed in Arg2^-/-^ striatum (98.48 ± 18.81 nmol/mg protein in Arg2^+/+^ vs. 194.61 ± 30.95 nmol/mg protein in Arg2-/-; p = 0.021), n = 7. **(C)** LC-MS-based measurements of combined polyamine levels and individual polyamine ratios in the striatum of Arg2+/+ and Arg2-/- mice. (Top left panel) No statistically significant differences in the combined Put+Spmd+Spm levels were observed between Arg2^+/+^ and Arg2^-/-^ striata (527.20 ± 82.04 nmol/mg protein in Arg2^+/+^ vs. 649.95 ± 79.70 nmol/mg protein in Arg2^-/-^; *p* = 0.304), n = 7. (Top middle panel) A statistically significant increase in the Spmd/Put ratio was observed in the Arg2^-/-^ striatum (8.03 ± 0.51 in Arg2^+/+^ vs. 14.20 ± 1.29 in Arg2^-/-^; *p* < 0.001), n = 7. (Top right panel) A statistically significant increase in the Spm/Spmd ratio was observed in the Arg2^-/-^ striatum (0.26 ± 0.021 in Arg2^+/+^ vs. 0.452 ± 0.043 in Arg2^-/-^; *p* = 0.0018), n = 7. (Bottom left panel) A statistically significant increase in the Spm/Put ratio was observed in the Arg2^-/-^ striatum (2.04 ± 0.14 in Arg2^+/+^ vs. 6.64 ± 1.10 in Arg2^-/-^; *p* = 0.0013), n = 7. (Bottom right panel) A statistically significant increase in the (Spmd+Spm)/Put ratio was observed in the Arg2^-/-^ striatum (10.07 ± 0.54 in Arg2^+/+^ vs. 20.84 ± 2.34 in Arg2^-/-^; *p* < 0.001), n = 7. **(D)** LC-MS-based measurements of N1-acetylspermidine (N1-AcSpmd) levels in the striatum of Arg2^+/+^ and Arg2^-/-^ mice. No statistically significant differences in the N1-AcSpmd levels were observed between Arg2^+/+^ and Arg2^-/-^ striata (10.89 ± 1.33 in Arg2+/+ vs. 8.38 ± 0.78 in Arg2-/-; p = 0.129), n = 7. Data are shown as mean ± SEM. *p < 0.05, **p < 0.01, ***p < 0.001, t-test.

We next examined enzymes involved in polyamine synthesis. We found that loss of Arg2 resulted in SRM overexpression (Fig. 9A), consistently with decreased Put and elevated Spmd/Put ratio in Arg2^-/-^ striata, suggesting that SRM upregulation drives excessive Put utilization. Despite increased Spm levels, SMS Unexpectedly, SMS levels remained unchanged (Fig. 9A), despite significant increase in Spm and lack of accumulation of Spmd in the presence of increased SRM expression. Immunohistochemistry confirmed SRM upregulation and showed its co-localization with DARPP-32 in MSNs (Fig. 9B). ODC, undetectable by western blot due used antibodies limitations, partially co-localized with DARPP-32 and showed significant alterations in Arg2^-/-^ striata: increased staining intensity, enhanced ODC-DARPP-32 co-localization and altered distribution with numerous longitudinal, ODC-positive clusters associated with DARPP-32-positive neurons (Fig. 9B).

**Fig. 9.**
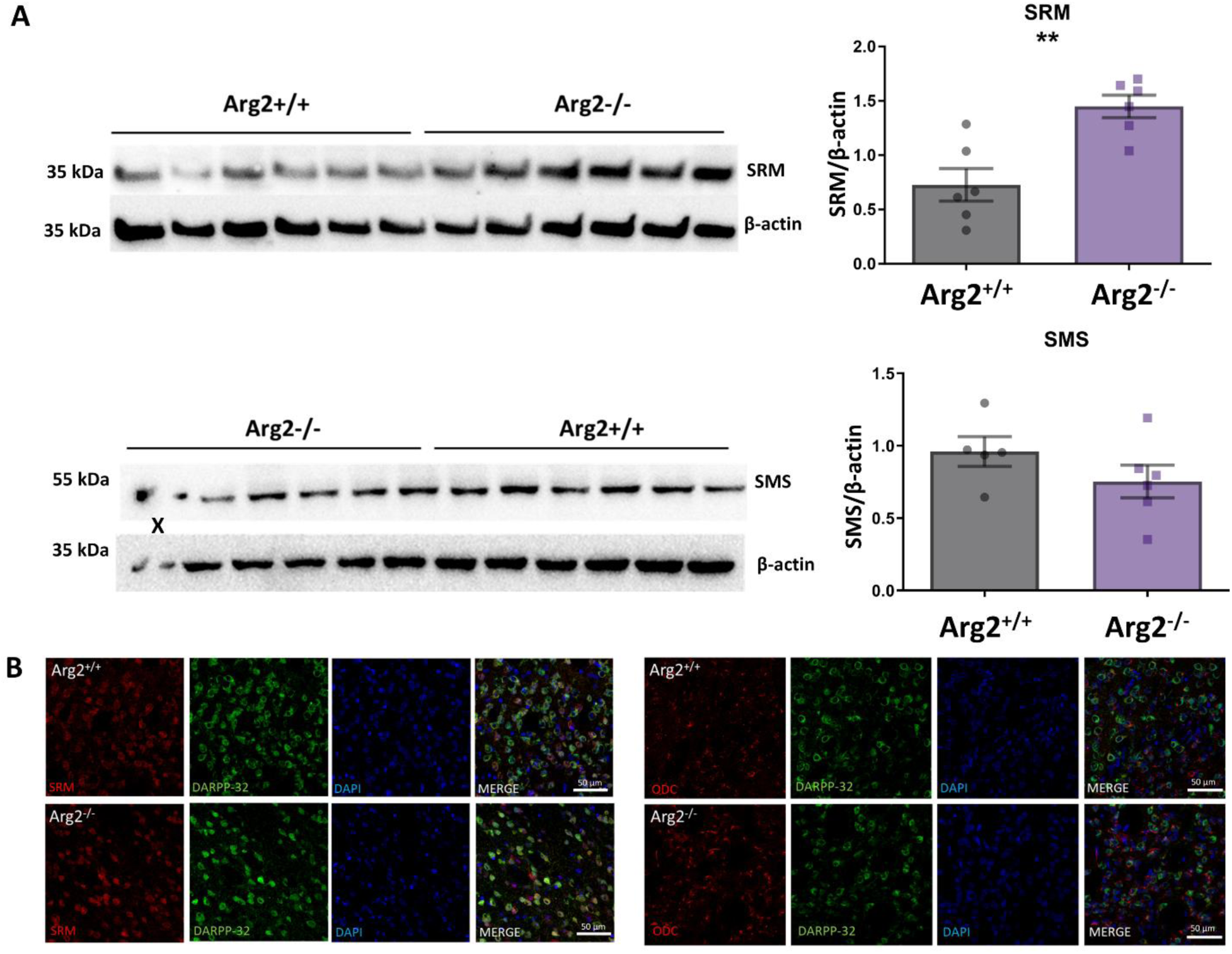
Analysis of spermidine synthase (SRM) and spermine synthase (SMS) expression and distribution in Arg2^+/+^ and Arg2^-/-^ mouse striata. **(A)** Measurements of SRM and SMS expression in Arg2^+/+^ and Arg2^-/-^ mouse striata. (Top left panel) Western blot image of SRM and β-actin in Arg2^+/+^ and Arg2^-/-^ striata. (Top right panel) A statistically significant increase in SRM expression was observed in Arg2^-/-^ striatum as determined by the quantification of SRM western blot normalized to β-actin western blot (0.73 ± 0.15 in Arg2^+/+^ vs 1.45 ± 0.10 in Arg2^-/-^ (arbitrary units); p = 0.003), n = 6. (Bottom left panel) Western blot image of SMS and β-actin in Arg2^+/+^ and Arg2^-/-^ striata. (Bottom right panel) No statistically significant differences in SMS expression were observed between Arg2^+/+^ and Arg2^-/-^ striata as determined by the quantification of SMS western blot normalized to β-actin western blot (0.96 ± 0.10 in Arg2^+/+^ vs 0.75 ± 0.11 in Arg2^-/-^ (arbitrary units); p = 0.217), n = 5-6. X indicates a sample identified as a technical outlier and excluded from the calculations. **(B)** Immunohistochemical analysis of SRM and ornithine decarboxylase (ODC) distribution in Arg2^+/+^ and Arg2^-/-^ mouse striata. (Left panel) Co-immunofluorescent staining of SRM (red) and the medium spiny neuron (MSNs) marker, DARPP32 (green), in the striata of Arg2^+/+^ and Arg2^-/-^ mice. Nuclei are counterstained with DAPI (blue). Representative images for Arg2^+/+^ and Arg2^-/-^ are shown. Scale bars (50 μm) are located in the right bottom corner of the merged images. SRM co-localized with DARPP-32 in the cell bodies of MSNs. SRM staining intensity was increased in Arg2^-/-^ striata. (Right panel) Co-immunofluorescent staining of ODC (red) and the MSNs marker, DARPP32 (green), in the striata of Arg2^+/+^ and Arg2^-/-^ mice. Nuclei are counterstained with DAPI (blue). Representative images for Arg2^+/+^ and Arg2^-/-^ are shown. Scale bars (50 μm) are located in the right bottom corner of the merged images. ODC partially co-localized with DARPP-32 in the cell bodies of MSNs. ODC staining intensity and ODC-DARPP-32 co-localization were increased in Arg2^-/-^ striata, along with an altered ODC distribution pattern, where ODC formed numerous longitudinal, intensely stained clusters associated with DARPP-32-positive neurons. Data are shown as mean ± SEM. **p < 0.01, t-test.

ODC, a key enzyme in polyamine synthesis, is tightly regulated, primarily by interaction with ornithine decarboxylase antizyme (OAZ; Fig. 10A), which targets ODC to proteasomal degradation. In Arg2^-/-^ striata, OAZ levels were unchanged compared to Arg2^+/+^ mice (Fig. 10B). OAZ-ODC interaction is modulated antizyme inhibitor (AZIN), which, by binding OAZ, prevents it from targeting ODC (Fig. 10A). We found a significant increase in AZIN1, brain-enriched AZIN isoform in Arg2^-/-^ striata (Fig. 10B), suggesting enhanced protection of ODC from degradation.

**Fig. 10.**
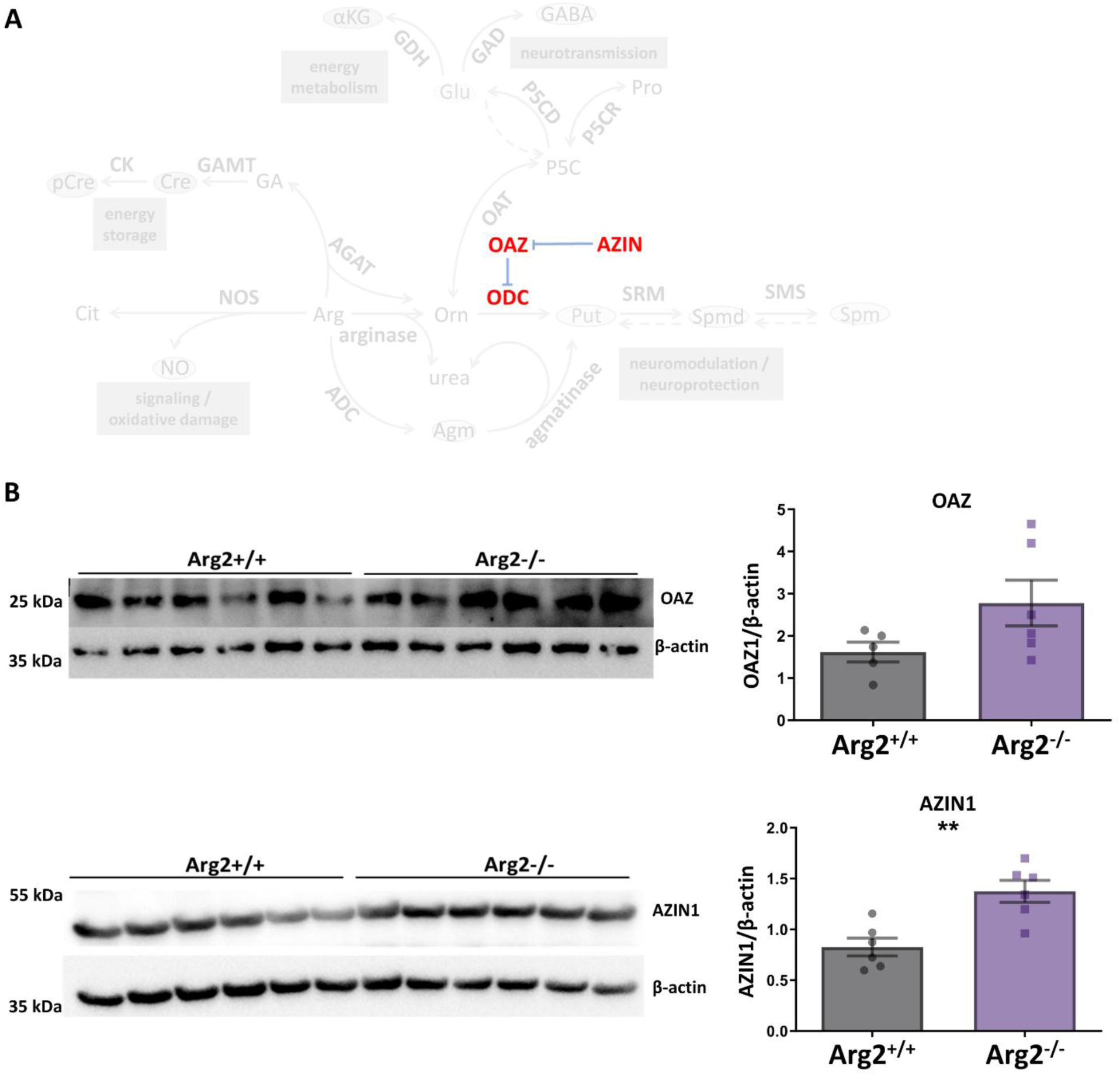
Analysis of ornithine decarboxylase (ODC)-regulating proteins in the striatum of Arg2^+/+^ and Arg2^-/-^ mice. **(A)** Schematic representation of the mechanism of ODC regulation by antizyme inhibitor (AZIN) and ornithine decarboxylase antizyme (OAZ). **(B)** Measurements of OAZ and AZIN1 expression in Arg2^+/+^ and Arg2^-/-^ mouse striata. (Top left panel) Western blot image of OAZ and β-actin in Arg2^+/+^ and Arg2^-/-^ striata. (Top right panel) No statistically significant differences in OAZ expression were observed between Arg2^+/+^ and Arg2^-/-^ striata as determined by the quantification of OAZ western blot normalized to β-actin western blot (1.61 ± 0.24 in Arg2^+/+^ vs 2.78 ± 0.54 in Arg2^-/-^ (arbitrary units); p = 0.101), n = 5-6. (Bottom left panel) Western blot image of AZIN1 and β-actin in Arg2^+/+^ and Arg2^-/-^ striata. (Bottom right panel) A statistically significant increase in AZIN1 expression was observed in Arg2^-/-^ striatum as determined by the quantification of AZIN1 western blot normalized to β-actin western blot (0.83 ± 0.09 in Arg2^+/+^ vs 1.37 ± 0.11 in Arg2^-/-^ (arbitrary units); p = 0.003), n = 6. Data are shown as mean ± SEM. **p < 0.01, t-test.

### Striatal Arg-related metabolite profile is altered in response to Arg2 deficiency

Finally, we employed principal component analysis (PCA) to visualize extended patterns in the metabolic profiles of polyamines and Arg2-related amino acids, based on our LC-MS data. PC1 and PC2 accounted for 52% and 26.86% of data variance, respectively (Fig. 11). The PC1 × PC2 score plot clearly separated Arg2^+/+^ and Arg2^-/-^ into distinct clusters, with greater dispersion within Arg2^-/-^ group (Fig. 11), indicating that Arg2 loss not only altered metabolite profile but also increased their variability.

**Fig. 11.**
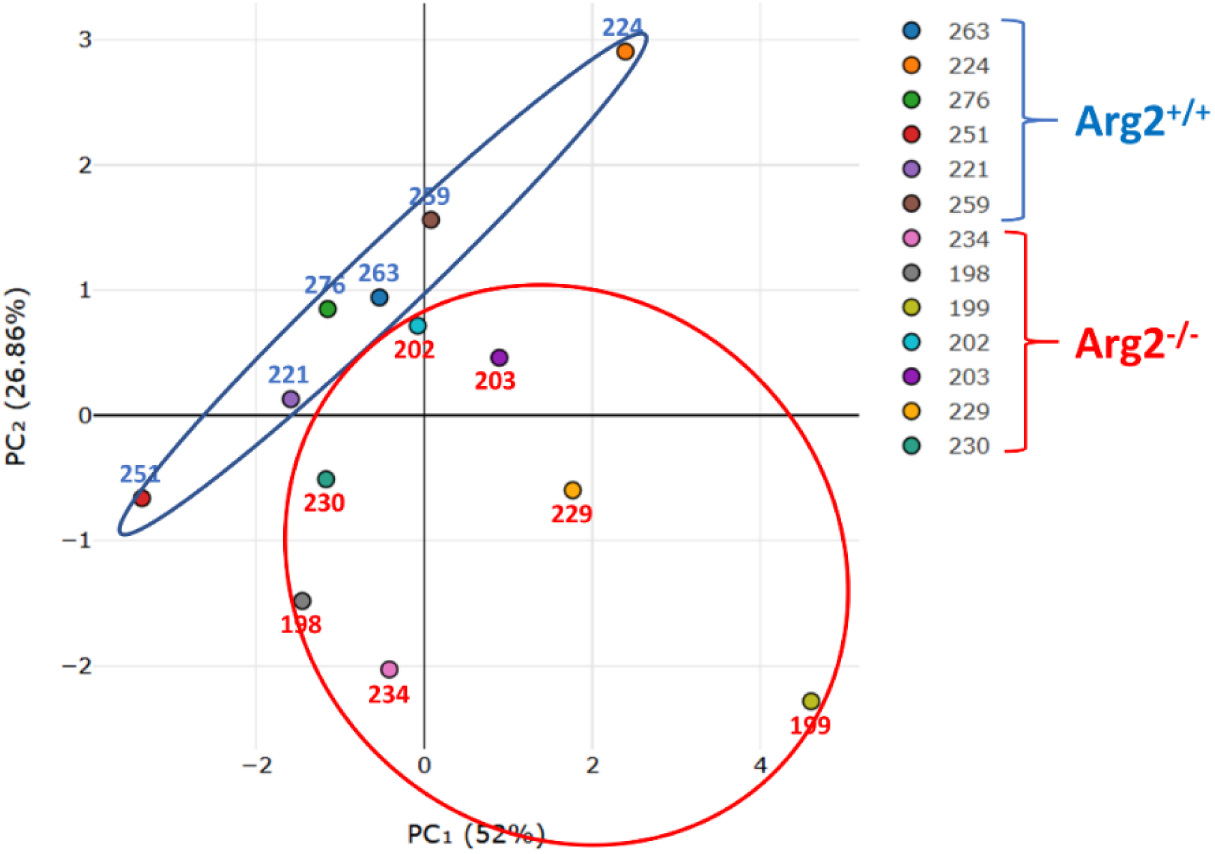
Multivariate Principal Component Analysis (PCA) of metabolic profiles of polyamines and Arg2-related amino acids in the striatum of Arg2^+/+^ and Arg2^-/-^ mice. The LC-MS dataset, which included putrescine, spermidine, spermine, N1-acetyl-spermidine, arginine, citrulline, ornithine and proline was used as input. Principal component 1 (PC1) and principal component 2 (PC2) accounted for 52% and 26.86% of the data variance, respectively. The PC1 × PC2 score plot separated Arg2^+/+^ and Arg2^-/-^ samples into two distinct clusters, encircled by blue and red ellipses, respectively, as shown in the graph.

## Discussion

To better understand Arg2 role in striatal Arg metabolism, we compared levels of Arg and related metabolites in Arg2^+/+^ and Arg2^-/-^ mice. As previously demonstrated, Arg2 is the only arginase isoenzyme present in the striatum and solely mediates arginase enzymatic activity [12,13]. Notably, Arg2 didn’t induce compensatory Arg1 expression [13], indicating that the observed metabolic changes in Arg2^-/-^ striata are not simply due to Arg2 elimination but rather total loss of physiological arginase activity.

### Arginine and Ornithine

A significant increase in Arg levels in Arg2^-/-^ striata was expected and was consistent with previous observations in plasma [14–16], lymphoid tissues [17], neural progenitor cells and whole-brain homogenates [18]. In whole-brain homogenates, Arg as measured by LC-MS, was elevated by ∼50%, while in striata, in our LC-MS assay, it increased by ∼120%, emphasizing striatal enrichment with Arg2. Despite this Arg accumulation, Orn levels remained paradoxically unaltered in Arg2^-/-^ striata, as confirmed by both LC-MS and HPLC. A non-significant trend towards increased Orn was observed, suggesting statistical significance was not reached due to large variations among Arg2^-/-^ individuals.

We identified two potential compensatory Orn sources in Arg2^-/-^ striata: AGAT-mediated Orn production from Arg and the Pro-Orn pathway. AGAT expression was significantly elevated, consistent with our previous findings implicating Cre pathway in striatal metabolic alterations following Arg2 loss [13]. Among the three metabolites of Cre pathway, only GA was increased [13](, suggesting selective AGAT upregulation in Arg2^-/-^ striata without full pathway activation. Although AGAT is mainly studied in Cre synthesis, elevated GA has been observed in the blood, cerebrospinal fluid (CSF), and post-mortem brain of Arg1-deficienct patients [19] and in plasma, liver, and kidneys of Arg1^+/-^ and Arg1^+/-^/Arg2^-/-^ mice [20], indicating AGAT activation in response to arginase loss. Similarly to our findings, [13], GA increase in blood was not accompanied by a Cre increase in most analyzed Arg1-deficient patients [21], suggesting that AGAT activation in attempt to support Orn production may be a common cellular response to arginase loss. In the brain, AGAT is localized primarily in glial cells, particularly oligodendrocytes [22], suggesting potential glia-MSNs interactions in regulating striatal Arg-to-Orn conversion.

We also found increased OAT expression, which catalyzes the reversible conversion between P5C and Orn. P5C can be produced from Pro by Pro oxidase and Pro-to-Orn pathway is prominent in neonates, however shifts towards Orn-to-Pro direction in adults [23,24]. In the adult rodent brain, Pro synthesis from Orn is well documented [25–27], while Pro-derived P5C is rather a precursor for Glu [28,29]. However, in Arg2^-/-^ striata, Orn/Pro ratio increased without changes in total Orn+Pro, suggesting a pathway shift towards Orn synthesis. This was supported further by increased Arg+Cit+Orn content, indicating alternative routes of Orn production beyond direct Arg-to-Orn conversion. Similarly to AGAT, OAT was localized outside MSNs – in DARPP-32-negative neuropil, again implying transcellular metabolic interaction to maintain striatal Orn homeostasis following Arg2 loss. While OAT can also participate in Glu to Orn conversion in some tissues [30–32], no differences in Glu or related metabolites (Gln, GABA, αKG) nor in Orn/Glu or Orn/(Glu+Gln+GABA) were observed between genotypes, suggesting this route is not relevant in the striatum.

The preservation of Orn levels seems to be a common cellular response to Arg2 loss across tissues. In Arg2^-/-^ mice, Orn remains unaltered in biofluids, cells or tissues, despite significant Arg accumulation [14–16,18]. Consistently, Arg2 overexpression under neuron-specific Thy1 promoter lowers Arg without affecting Orn in the brain [33], whereas Thy1-driven Arg1 overexpression increases Orn, suggesting the two isoenzymes play distinct roles in Orn homeostasis. This differences may reflect existence of distinct subcellular Arg and/or Orn pools, with Arg2-linked pool being closely monitored and buffered by compensatory mechanisms, unlike Arg1-related pool. In this context, the previously mentioned AGAT upregulation in Arg1 deficiency [19,20] might be insufficient to fully normalize Orn levels, further supporting this distinction.

In summary, Orn homeostasis in the striatum appears to be maintained by a dynamic balance of multiple intersecting pathways. Despite marker Arg increase in Arg2^-/-^ tissue, Orn levels are restored via alternative metabolic pathways, including AGAT and OAT upregulation, facilitating alternative Arg utilization and Pro-to-Orn conversion, respectively.

### Nitric oxide

An important finding of our study is the lack of changes in NOx levels in Arg2^-/-^ striatum. Similarly, Cit levels – the second product of NOS activity – remained unaltered. Since Orn transcarbamylase, the only other Cit-producing enzyme, is absent in the healthy brain [34,35], Cit can serve as an indicator of NO production [36]. Additionally, nNOS expression, the major NOS isoenzyme in the non-diseased brain [37], was unchanged. Thus, excessive Arg accumulation following Arg2 loss doesn’t affect activity or expression of nNOS, indicating that Arg2 is not involved in regulating striatal NO synthesis.

Though arginase and NOS both utilize Arg, their competition remains controversial Arginase has ∼1000-fold higher Km for Arg than NOS [38,39], but also ∼1000-fold higher Vmax [39,40], what theoretically allows them to compete equally. This has been shown in vitro in cells endogenously co-expressing both enzymes, like endothelial cells [41,42] and stimulated macrophages [43,44], where arginase inhibition increased NO production. However, in vivo relevance is debated due to potential existence of distinct, poorly interchangeable Arg pools, as shown by existence of “arginine paradox”, where exogenous Arg stimulates NO production despite saturating endogenous Arg levels, suggesting that NOS has access only to a fraction of the total cellular Arg [44–47]. This effect may be particularly relevant to Arg2, which is mitochondrial, whereas NOS isoforms are cytosolic. Moreover, most studies supporting competition were performed under Arg-limited conditions, whereas *in vivo* cells continuously import Arg from the extracellular space. Recent study demonstrated that constant extracellular Arg supply prevents arginase-mediated inhibition of NO production [48], suggesting minimal competition under physiological conditions *in vivo*.

In the striatum, inter-cellular Arg2 and NOS compartmentalization may further limit potential competition for Arg. Arg2 is expressed in MSNs [13], while nNOS is localized mainly in nitrergic interneurons [49,50], and the second physiological NOS isoform, endothelial NOS (eNOS) - in endothelial cells. Consequently, Arg pools available to Arg2 and NOS are separated by at least three barriers: MSNs mitochondrial membrane, MSNs cellular membrane, nitrergic neuron or endothelial cell cellular membrane. Arg transport, therefore, becomes a key for the involvement of both enzymes in metabolism of striatal Arg, as it was shown in the study, where inhibition of y^+^LAT2 Arg transporting system reduced striatal NO synthesis [51]. Additionally, considering successful AGAT upregulation in Arg2^-/-^ striatum, one can hypothesize that a fraction of Arg may be driven to glial cells for Orn production, potentially limiting its availability for NO synthesis.

Our conclusion apply to physiological conditions, but Arg2 loss still may impact NO synthesis in various pathologies, where NO production increases significantly. Such an effect was found in macrophages, where *H. Pylori* infection resulted in greater NO production and iNOS expression in Arg2^-/-^, while in non-infected cells, similarly to our findings, no difference in NO synthesis were observed between Arg2^-/-^ and Arg2^+/+^ mice. [52]. A similar mechanism could also apply in the striatum during pathology, like PD or HD, disorders, where neuroinflammation is a crucial step the pathogenesis. HD is associated with both, elevated striatal iNOS expression [53], and progressive striatal Arg2 impairment [54]. This could be a mechanism contributing to increased susceptibility of MSNs, a neuronal population that specifically expresses Arg2 [13], to injury in HD, although further experimental confirmation is required.

### Polyamines

Arginases regulate polyamine production by supplying Orn for the first step of the pathway: ODC-mediated conversion of Orn to Put. In the brain, this step is critical because polyamines only minimally cross the blood-brain barrier [55,56]. Unlike other tissues, the brain can’t rely on circulating polyamines, thus the availability of Arg and Orn, along with the expression and activity of polyamine-related enzymes and their regulatory proteins, is essential for cerebral polyamine homeostasis.

We demonstrate that the total striatal polyamine content (Put+Spmd+Spm) remains unaltered in Arg2^-/-^ mice, likely due to compensation through alternative Orn-generating pathways, most likely AGAT- and/or OAT-dependent. The potential involvement of both in polyamine synthesis has been demonstrated in other tissues. Elevated AGAT mRNA expression correlated with polyamine accumulation in splenic myeloid cells from glioma-bearing mice [57], while OAT was shown to support polyamine synthesis in endothelial cells [58]. OAT was also found to be a major Orn supplier in pancreatic cancer [59], whereas in healthy pancreas, arginase-derived Orn dominated [60]. In these reports, however, Gln was identified as a major source of P5C. A role for Pro is best described in pigs, where it was shown that Pro can serve as a source of polyamines in porcine placenta [61] and intestines of cortisol-exposed piglets [62,63]. These examples confirm the existence of AGAT- and OAT-mediated pathways supporting polyamine synthesis, whose significance varies depending on species, tissue type and pathological states.

Although total polyamines were unchanged, two of the three differed between groups: Put decreased, while Spm increased in Arg2^-/-^ striata, resulting in a significant increase in higher-to-lower polyamine ratios (Spmd/Put, Spm/Spmd, Spm/Put), indicating a shift towards higher polyamines. This was accompanied by altered expression of certain enzymes and regulating proteins. AZIN1 was upregulated, suggesting higher ODC stability through inhibition of OAZ-mediated degradation. Consistently, ODC staining was stronger in Arg2^-/-^ striata, although the substrate (Orn) and product (Put) levels didn’t reflect elevated ODC expression. Orn remained unchanged possibly due to sustained supply from AGAT and OAT and/or feedback inhibition by significantly increased Spm, the strongest endogenous ODC inhibitor [64]. Put depletion, despite stable ODC, may result from increased SRM expression, which accelerates its conversion to Spmd. Notably, Spmd levels remained unchanged, suggesting efficient conversion to Spm pool, consistent with exceptionally high SMS activity in the brain, the only tissue where it exceeds that of SRM [65].

These data indicate that while Arg2 is not essential for polyamine synthesis in the striatum, its absence significantly disrupts their balance. This occurs despite maintained Orn levels, suggesting that not only Orn availability but also its source affects individual steps of Orn-to-Spm pathway. Additionally, metabolic signals resulting from Arg2 loss (e.g. massive Arg accumulation) may trigger additional pathways contributing to the observed imbalance in striatal polyamines and related amino acids, as reflected in distinct Arg-related metabolic profiles of Arg2^+/+^ and Arg2^-/-^ striata detected by PCA. Overall, our findings highlight the key role of Arg2 in supporting polyamine balance by maintaining striatum-specific Arg-to-Orn conversion rate. The exceptional enrichment of Arg2 in the striatum [13,54] suggests that Arg2-dependent polyamine homeostasis is essential for certain striatal functions, particularly in MSNs, where Arg2 is localized.

The importance of Arg2 for polyamine homeostasis is established in some peripheral tissues. In liver and small intestine, Arg2^-/-^ mice exhibited ∼4-fold increase in Put with unaffected Spmd and Spm levels [66], while in endothelial cells, Arg2 overexpression increased Put and Spmd [67]. In Arg2^-/-^ retina, however, polyamine levels were unchanged [68], similarly to whole-brain homogenates [66], highlighting striatal unique sensitivity to Arg2 loss.

Polyamine dysregulation is implicated in many brain disorders. In a mouse model of hypoxia-ischemia, Spmd and Spm decreased in affected area [69], whereas in rat model of traumatic brain injury (TBI), Put increased [70,71]. The relevance of these changes to disease remains unclear, as interventions in polyamines yield mixed results: exogenous Spm protected against hypoxic injury in rat neonates [72], whereas ODC inhibition reduced infarct volumes in stroke [73] and improved neurological outcomes in TBI [71] in rat models.

In neurodegenerative diseases, polyamine upregulation was reported in disease-affected regions in Alzheimer’s disease (AD) patients [74] and in transgenic AD mouse models [75,76]. ODC overexpression impaired memory [77], while ODC knockdown cleared amyloid β (Aβ) plaques and shifted astrocytes to a regenerative state in APP/PS1 AD mice [78]. Difluoromethylornithine (DFMO), an ODC inhibitor, reduced Aβ deposition and improved cognition function in another AD model [79], leading to a single-patient clinical trial, in which, however, DFMO failed to suppress cognitive decline and amyloidosis progress in mild cognitive impairment patient, and didn’t prevent the progress to AD during this on-year trial [80]. The role of polyamines in AD/dementia progression is not unequivocally negative. DFMO was also shown, in contrast to the above data, to impair learning [81], while Spmd improved memory in rats [82] and enhanced cognition in patients at risk of dementia [83]. These contradictory findings highlight the complex roles of polyamines, suggesting that maintaining their region-specific balance, not only the total levels is crucial for brain health

## Methods

### Materials and reagents

Acetonitrile LC-MS grade (J.T. Baker); Acetonitrile, UPLC grade (J.T. Baker); acryl:bis (VWR); Agmatine sulfate salt (Sigma-Aldrich); L-Arginine (Sigma-Aldrich); L-Arginine-5-13C,4,4,5,5-d4 (Sigma-Aldrich); BCA Protein Assay Kit (Sigma Aldrich); bromophenol blue (BPB) (Sigma-Aldrich); centrifuge filters (Amicon); L-Citrulline (Sigma-Aldrich); DAPI (ITW Reagents); EtOH (Stanlab); Fluor Save Reagent (Millipore); Formic acid (Sigma-Aldrich); glycerol (Chempur); Hypersil ODS C18 column (Sigma-Aldrich); Methanol, LC-MS grade (J.T. Baker); milk (Gostyn); Milli-Q-grade water (Millipore); Mouse on Mouse (M.O.M) kit Blocking Reagent (Thermo-Fisher Scientific); N1-Acetylspermidine Dihydrochloride (TRC); N1 Acetylspermidine-d3 Dihydrochloride (TRC); N1-acetylspermine Trihydrochloride (TRC); N1-acetylspermine-d3 Trihydrochloride (TRC); NaCl (Chem-Pur); NaOH (Pol-Aura); Nitrate/Nitrite Fluorometric Assay Kit (Cayman Chamicals); NP-40 (Santa-Cruz); o-phthaldialdehyde (OPA) (Sigma-Aldrich); L-Ornithine hydrochloride (Sigma-Aldrich); L-Ornithine-13C5 hydrochloride (Sigma-Aldrich); Paraformaldehyde (PFA) (Lach-Ner); Pentobarbital (Biowet, Poland); Phosphate-buffered saline (PBS) (Gibco); PolyFreeze Tissue Freezing Medium (Sigma-Aldrich); Ponceau S (Pol-Aura); L-Proline (Sigma-Aldrich); Propionic Anhydride (Sigma-Aldrich); protease inhibitor cocktail (Roche); Putrescine-d8 (TRC); Pyridine (Sigma-Aldrich); SDS (Chempur); sodium azide (VWR); Sodium bicarbonate (Sigma Aldrich); sodium citrate (Sigma-Aldrich); Spermidine trihydrochloride (TRC); Spermidine-d6 Trihydrochloride (TRC); Spermine tetrahydrochloride (Sigma-Aldrich); Spermine-(butyl-d8) tetrahydrochloride (Sigma-Aldrich); Standards to High-Performance Liquid Chromatography (HPLC) measurement: Asn, Ser, His, Thr, Cit, Arg, Ala, Trp, Val, Phe, Ile, Leu, Orn, Asp, Glu, Gln, Tau, GABA (Sigma-Aldrich); sucrose (Lach-Ner); SuperSignal West Dura (Thermo Scientific); TEMED(Alfa-Aesar); Tris (Chem-pur); Tris base (Chem-pur) ; Triton x-100 (Sigma-Aldrich); Tween-20 (Sigma-Aldrich); Teflon-glass homogenizer (Kimble); α-Ketoglutarate Assay Kit (Sigma-Aldrich); β-mercaptoethanol (β-Me) Acros Organics)

### Animals

Arg2-deficient (Arg2^tm1Weo^/J strain; Jackson Laboratory; hereafter referred to as Arg2^-/-^) and wild-type control (C57Bl/6J; hereafter referred to as Arg2^+/+^) mice were used. Animals were housed in the Animal Facility of Mossakowski Medical Research Institute (MMRI PAS) under a 12-hour light/dark cycle, with food and water available *ad libitum*. Procedures were performed under approved licenses and were designed to minimize pain and distress. The study complied with EU Directive 2010/63/EU and Polish legislation (Act no. 266/15.01.2015). Gender-balanced groups ∼3 month-old animals were used.

### Tissue preparation

Tissue was prepared as described in [13] with modifications. For metabolite analysis, animals were sacrificed, and the brains were rapidly removed, frozen, and stored at −80°C. Striata were dissected from frozen brain slices on wet ice under SMZ800 N stereomicroscope (Nikon, Japan), pooled, weighed and homogenized in an assay-appropriate buffer using a Teflon-glass homogenizer. Homogenates were then processed according to assay requirements. For histology, animals were deeply anesthetized via with pentobarbital and perfused with PBS (pH 7.4) followed by 4% paraformaldehyde (PFA). Brains were post-fixed in 4% PFA for 24 hours at 4°C, cryoprotected in 30% sucrose with 0.1% sodium azide and embedded in PolyFreeze. Frozen brains were sectioned at 30 μm using a Hyrax M 25 microtome (Zeiss, Germany) with an MTR freezing unit (SLEE Medical, Germany) at −35°C. The slices were stored in PBS with 0.1% sodium azide.

### Protein content measurement

Total protein content was measured for selected assays using the BCA Kit. Samples were transferred to 96-well plates, mixed with BCA reagent supplemented with 0.08% CuSO₄. And incubated for 30 min. at 37°C. Absorbance was measured at 562 nm using the Infinite M1000 PRO plate reader (Tecan, Switzerland). Protein concentrations were calculated from a standard curve run on the same plate.

### Liquid chromatography - mass spectroscopy

Arg, related amino acids, and polyamines were measured using liquid chromatography-mass spectrometry (LC-MS). Striatal tissue was homogenized on ice in 70% ethanol (EtOH; 90 μl/10 mg), centrifuged at 14 000 × g for 15 min. at 4°C (Mikro 220R centrifuge; Hettich, Germany). Pellets were dissolved in 60 μl of 0.1 M NaOH with 0.125% Triton X-100 for protein measurements and 10 μl of supernatants were mixed in low-retention tube with 10 μl of 0.5 M sodium bicarbonate and 50 μl of internal standard (putrescine-d8, spermine-d8, spermidine-d6, arginine-d5, ornithine-d5, N1-acetylspermidine-d3, and N1-acetylspermine-d3) dissolved in HPLC-grade acetonitrile. Samples were then vortexed (1 min., 1 500 RPM), 100 μl of derivatization mixture (propionic anhydride/acetonitrile/pyridine, 1:9:0.2, v/v/v) was added, samples were vortexed again (45 min., 800 RPM, 50°C), evaporated under nitrogen and reconstituted in 200 μl of 10% acetonitrile with 0.1% formic acid. Alongside, a 9-point calibration curve was prepared for each analyte in 70% EtOH.

Metabolite profiling was performed using an Acquity UPLC system (Waters, USA) with a Xevo TQ-S mass spectrometer (Waters). Separation was performed using an ACQUITY UPLC BEH C18 column (1.7 μm, 2.1 mm × 100 mm; Waters). Mobile phase A consisted of Milli-Q water with 0.1% formic acid, and mobile phase B of methanol (MetOH)/acetonitrile (3:7). Flow rate was 0.5 ml/min., and running time was 4.5 min. The spectrometer operated in positive electrospray ionization mode, and mass spectra were acquired in multiple-reaction-monitoring mode. Data were analysed using MassLynx V4.2 (Waters) and normalized to protein content.

### Nitrates and nitrites assay

Tissue was homogenized in 100 μl PBS and centrifuged at 18 620 x g for 20 min. at 4°C in Mikro 220R centrifuge. A portion was reserved for protein measurement and the rest was ultracentrifuged at 120 000 x g for 30 min. at 4°C (Optima MAX-XP ultracentrifuge; Beckman, USA). Supernatants were filtered through 10 kDa centrifuge filters to remove contaminants. NOx concentrations were measured using a NOx Kit. Nitrates were converted to nitrites by 2 h incubation with nitrate reductase at 21°C, followed by 10 min. reaction with 2,3-diaminonaphthalene (DAN) at room temperature (RT) to form fluorescent 1(H)-naphthotriazole. NaOH was added to enhance fluorescence, and samples were measured at λex 360 nm and λem 430 nm in white 96-well plate using an Infinite M1000 PRO reader. NOx concentrations were determined from a standard curve generated on the same plate and were normalized to the protein content.

### High-performance liquid chromatography

A panel of 18 amino acids was analyzed in striatal tissue of Arg2^+/+^ and Arg2^-/-^ mice by HPLC, as described [84,85] using an UltiMate 3000 chromatograph (Thermo Scientific, USA). Tissue was homogenized on ice in 70 μl PBS and centrifuged at 18 620 x g for 15 min. at 4°C (Mikro 220R). Supernatants were mixed with MetOH (1:9) and deproteinized by centrifugation. Samples were derivatized with 1.5 mM OPA in borate buffer and 60 μl was injected onto a Hypersil ODS C18 column (5 μm, 4.6 mm x 250 mm). Asn, Ser, His, Thr, Cit, Arg, Ala, Trp, Val, Phe, Ile, Leu and Orn were separated with 50 mM acetate buffer (pH 6.8, flow rate: 1.5 ml/min). Asp, Glu, Gln, Tau, and GABA were separated with 50 mM phosphate (pH 6.2) + 10% MetOH (flow rate: 1.2 ml/min). In both cases, 100% MetOH served as mobile phase B, with 20%-70% gradient over 30 min.

Detection was performed using an RF 2000 fluorescence detector (Dionex, USA, λex 340 nm, λem 450 nm). Concentrations were calculated from peak areas of standard curves of OPA-derivatized amino acids. Data acquisition and analysis were performed using Chromeleon 6.8 (Dionex, USA). The results were normalized to tissue mass.

### α-Ketoglutarate measurements

The levels of αKG were measured using fluorimetric αKG Assay Kit. Tissues were homogenized on ice in 100 μl of Assay Buffer, centrifuged at 13 000 x g for 10 min. at 4°C (Mikro 220R) and supernatants were filtered through 10 kDa centrifuge filters (Mikro 220R). 50 μl of sample was mixed with 50 μl of the Reaction Mix (assay buffer, converting enzyme, development enzyme mix and fluorescent peroxidase substrate), and incubated at 37°C for 30 min. Fluorescence (λex 535, λem 587 nm) was measured with an Infinite M1000 PRO reader, αKG concentrations were calculated from a standard curve run on the same plate and normalized to tissue weight.

### Western blotting

Enzymes from Arg-related pathways were measured by western blot as described in [13]. Tissue was homogenized on ice in 80 µl RIPA buffer (50 mM Tris, pH 8.0, 150 mM NaCl, 0.1% SDS, 0.05% NaDC, 1% NP-40) with protease inhibitors and centrifuged at 14 000 x g for 20 min. at 4°C (Mikro 220R). Following equalization of protein concentrations, samples were mixed (2:1) with Laemmli buffer (30% glycerol, 0.187 M Tris, 0.033% BPB, 15% β-Me), heated (95°C, 10 min.), and loaded (40 µg protein/well) onto polyacrylamide gel (4% stacker (0.12 M Tris, 4% acryl:bis, 0.1% SDS, 0.05% APS, 0.12% TEMED, pH 6.8) and 12% resolver (0.37 M Tris, 12% acryl:bis, 0.1% SDS, 0.05% APS, 0.05% TEMED, pH 8.8)). Electrophoresis was performed at 120 V for 2 h (Mini-Protean; Bio-Rad, USA) in a running buffer (191 mM Gly, 24.7 mM Tris base, 0.1% SDS) and proteins were transferred (Mini Trans-Blot Module; Bio-Rad) to nitrocellulose membrane (Amersham, Germany) at 0.3 A for 2 h in transfer buffer (25 mM Tris, 200 mM Gly, 10% MetOH). Transfer was assessed by Ponceau S (1.3 M in 832 mM CH3COOH) staining. Membranes were blocked with 5% milk in TBST (20 mM Tris, 150 mM NaCl, 0.1% Tween-20), incubated overnight at 16°C with primary antibodies (Table 1), and 2 h at RT with peroxidase-conjugated secondary antibodies (Table 1). Bands were visualized using SuperSignal West Dura in the FusionX imager (Vilber, France). Membranes were stripped with stripping buffer (2% SDS, 0.5 M Tris, 0.008% β-Me, pH 6.8) and re-probed for β-actin. Signal intensity was quantified using Fiji (NHS, USA) and normalized to β-actin.

**Table 1.** List of used antibodies. WB – concentrations used for western blotting. IF – concentrations used for immunofluorescence.

| Antibody | company | cat. no. | host | WB | IF |
| --- | --- | --- | --- | --- | --- |
| AGAT (GATM) | Abclonal | A6598 | rabbit | 1 : 1000 | 1 : 1 000 |
| Alexa Fluor-conjugated (488 nm) 2' Ab | Invitrogen | A11017 | mouse |  | 1 : 2 000 |
| Alexa Fluor-conjugated (555 nm) 2' Ab | Invitrogen | A21430 | rabbit |  | 1 : 2 000 |
| AZIN1 | Abclonal | A4747 | rabbit | 1 : 1000 | 1 : 250 |
| DARPP-32 | Santa Cruz | sc-271111 | mouse |  | 1 : 1 000 |
| HRP conjugated 2' Ab | Sigma-Aldrich | A9169 | rabbit | 1 : 2000 |  |
| HRP conjugated 2' Ab | Sigma-Aldrich | A9044 | mouse | 1 : 2000 |  |
| nNOS | BD Transduction | 610308 | mouse | 1 : 1000 |  |
| OAT | Abclonal | A6235 | rabbit | 1 : 1000 | 1 : 250 |
| OAZ | Abclonal | A744 | rabbit | 1 : 1000 |  |
| ODC | Abclonal | A1948 | rabbit |  | 1 : 250 |
| SMS | OriGene | TA503099 | mouse | 1 : 1000 |  |
| SRM | Abclonal | A8151 | rabbit | 1 : 1000 | 1 : 250 |
| β-actin | Abcam | ab49900 | mouse | 1 : 50 000 | |

### Immunostaining

Fluorescent immunostaining was performed as described in [13] with minor modification. Tissue slices were washed in PBS, and subjected to antigen retrieval in 10 mM sodium citrate (pH 6.0, 80°C, 30 min.). Non-specific binding was blocked with M.O.M Blocking Reagent (1:1000) in PBST (PBS + 0.5% Tween-20) followed by overnight incubation at 4°C with mouse anti-DARPP-32 (1:1000) and rabbit anti-Arg related enzyme (see Table 1) antibodies diluted in M.O.M Protein Concentrate in PBST (1:800). Slices were next incubated for 3 h at RT with Alexa Fluor-conjugated secondary antibodies (488 nm anti-mouse, 555 nm anti-rabbit) in 5% NHS in PBST. After DAPI counterstaining (1: 10 000, 10 min., RT), slices were mounted and coverslipped with Fluor Save. Imaging was performed using an LSM 780/ELYRA PS.1 confocal microscope (Zeiss) at the Laboratory of Advanced Microscopy Techniques, MMRI PAS, and data were analyzed using Zen 3.5 (Zeiss).

### Statistical analysis

Univariate comparisons between Arg2^+/+^ and Arg2^-/-^ mice were performed using unpaired t-test in GraphPad Prism 10.0.0 (GraphPad Software, USA). p-values less than 0.05 were considered significant. Data are presented as mean ± SEM.

Multivariate analysis of LC-MS data was performed by principal component analysis (PCA) using Statistics Kingdom tool (https://www.statskingdom.com). The analysis was based on the correlation matrix of standardized variables, and visualized as a 2D scatter plot (PC1 vs. PC2).

## Conclusions

In contrast to most brain regions where Arg2 expression is low or absent, striatal Arg2 plays a central role in regulating Arg-related metabolism, ensuring proper Arg utilization and maintaining the polyamine balance required for striatum-specific functions. Arg2 loss disrupts striatal Arg metabolism, leading to widespread changes in Arg-related metabolites and proteins – likely as both compensatory mechanisms and pathological consequences. This may trigger further consequences, such as the impairment of more distant metabolic pathways not directly related to Arg, as we observed before [13], what may consequently lead to striatal dysfunction. Therefore, Arg2 may represent a novel factor in striatum-specific pathologies, such as HD, where its impairment appears at presymptomatic stage and progresses with disease, although its significance in disease pathogenesis remains unknown.

## Acknowledgements

This work was supported by National Science Centre, Poland (grant number 2018/30/E/NZ1/00144) (**MW**) and by statutory funds no 13 from Mossakowski Medical Research Institute, Polish Academy of Sciences (**MZ**). During the preparation of this manuscript, authors used OpenAI ChatGPT 4.0 in order to improve language and readability. After using this tool, the authors thoroughly reviewed and edited the manuscript and take full responsibility for its content.

## Author contribution

**MN:** Conceptualization, Formal analysis, Investigation, Methodology, Project administration, Supervision, Validation, Visualization, Writing – original draft, Writing – review & editing, **OB:** Investigation, **MR:** Investigation, Methodology, Validation, Writing – review & editing, **KS:** Investigation, Methodology, **AS:** Investigation, Supervision, Writing – review & editing, **AO:** Investigation, **AD:** Investigation, **WH:** Methodology, Supervision, Validation, Writing – review & editing**, MZ:** Funding acquisition, Resources, Supervision, Writing – review & editing, **ES:** Methodology, Resources, Supervision, Writing – review & editing, **MW:** Conceptualization, Formal analysis, Funding acquisition, Methodology, Project administration, Resources, Supervision, Validation, Visualization, Writing – original draft, Writing – review & editing

